# A mechanism for migrating bacterial populations to non-genetically adapt to new environments

**DOI:** 10.1101/2021.09.21.461202

**Authors:** Henry Mattingly, Thierry Emonet

## Abstract

Populations of chemotactic bacteria can rapidly expand into new territory by consuming and chasing an attractant cue in the environment, increasing the population’s overall growth in nutrient-rich environments. Although the migrating fronts driving this expansion contain cells of multiple swimming phenotypes, the consequences of non-genetic diversity for population expansion are unknown. Here, through theory and simulations, we predict that expanding populations non-genetically adapt their phenotype composition to migrate effectively through multiple physical environments. Swimming phenotypes in the migrating front are spatially sorted by chemotactic performance, but the mapping from phenotype to performance depends on the environment. Therefore, phenotypes that perform poorly localize to the back of the group, causing them to selectively fall behind. Over cell divisions, the group composition dynamically enriches for high-performers, enhancing migration speed and overall growth. Furthermore, non-genetic inheritance controls a trade-off between large composition shifts and slow responsiveness to new environments, enabling a diverse population to out-perform a non-diverse one in varying environments. These results demonstrate that phenotypic diversity and collective behavior can synergize to produce emergent functionalities. Non-genetic inheritance may generically enable bacterial populations to transiently adapt to new situations without mutations, emphasizing that genotype-to-phenotype mappings are dynamic and context-dependent.

## Introduction

Groups of bacteria can migrate collectively by consuming an attractant cue in their environment, creating a traveling gradient that they chase (1–19). In extended environments, this process leads to directed range expansion and can dramatically increase the population’s overall growth, relative to undirected expansion, by putting cells in contact with nutrients before they are locally depleted (14,20). Even isogenic migrating populations contain multiple swimming phenotypes (12,21)—quantified, for example, by the fraction of time a cell spends tumbling, or tumble bias (22,23). However, the costs and benefits of phenotypic diversity for expanding bacterial populations are unknown.

Non-genetic diversity may be useful to migrating populations that encounter changing contexts, such different environments. A growing body of work has shown that the mapping from swimming phenotype to chemotactic performance—i.e. how fast the cell climbs a gradient—depends on the physical environment in which the cells swim. In liquid, cells with low tumble bias climb static attractant gradients the fastest (12,24) because they spend the least time reorienting and because their long runs allow them to more strongly bias their motion up the gradient. In porous environments like soil or semisolid agar, more frequent reorientations are needed to avoid obstacles in the media (4,10,11,25). However, it is unclear how a population can leverage the different strengths of its individual phenotypes during migration.

Our recent work may provide a clue. Experiments and quantitative modeling revealed that differences in chemotaxis abilities among individuals in the group are compensated by a leader-follower structure (12): high-performing phenotypes localize to the front of the group, where the gradient signal is weakest, and low-performing phenotypes localize to the back of the group, where the gradient signal is strongest. This sorting mechanism equalizes chemotactic drift speed throughout the group, allowing multiple phenotypes to travel together. But cells’ finite sensitivity for the attractant they chase causes a slow loss of cells at the back of the traveling group, where the attractant concentration gets small for the cells to detect the gradient. Furthermore, by placing certain phenotypes at the back of the group, spatial organization may cause some phenotypes to fall behind faster than others.

This slow loss of cells must be balanced by growth to maintain migration, but growth with multiple phenotypes requires considering which phenotypes are produced when a cell divides. In particular, to what extent do daughter cells non-genetically inherit the phenotypes of their mothers? Recent experiments have demonstrated non-genetic inheritance of swimming behaviors (12,26) and measured the correlation between mother and daughter phenotypes (26). But it is unclear how this inheritance might contribute to the phenotypic composition, migration, and growth of traveling populations.

Here, using simulations and analytical theory, we predict that an isogenic population can non-genetically adapt its phenotype composition to migrate effectively through multiple environments. By placing low-performers at the back of the group and at highest risk of falling behind, collective migration selectively removes those phenotypes. To understand this, we derive approximate analytical expressions for how the leakage flux of each phenotype depends on group properties in a generalized Keller-Segel model (2). Cell growth and division imperfectly replace the lost phenotypes, depending on non-genetic phenotypic inheritance. As a result, the long-term group composition dynamically and stably shifts towards high-performing phenotypes, increasing the group’s migration speed and the entire population’s growth rate. Analytically deriving the dependence of steady state migration speed on group properties, we find that the lowest performers in the group determine the group’s traveling speed. Finally, we show that in rapidly-changing environments, diversity becomes a liability to migration; but in slowly-changing ones, an optimal level of phenotypic inheritance allows a diverse population to out-run a non-diverse population. This is achieved by balancing a trade-off between large shifts in phenotypic composition and fast adaptation to new environments. More broadly, these results show that collective behaviors exert a selective pressure on the performance of individuals in the population, which, with growth, dynamically adapts its composition to the demands of the collective task. Thus, collective behavior and phenotypic diversity can synergize to provide an isogenic population with emergent functionalities.

## Results

### Spatial organization in the migrating group is environment-dependent

Past work has demonstrated that non-genetic diversity can benefit populations when individuals are better adapted to different situations (27–45). Therefore, we reasoned that diverse swimming phenotypes may benefit a collectively-migrating population if they are better at performing chemotaxis in different contexts. To study this quantitatively, we constructed a mathematical model of collective migration that extends the classic Keller-Segel model (2) in two ways (12). First, the model includes multiple swimming phenotypes, and second, it accounts for cells’ finite sensitivity for the attractant. Without growth, the model consists of the following partial differential equations:

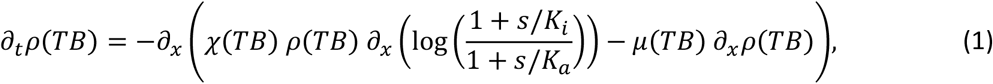

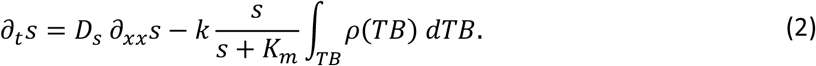

Here, *ρ*(*TB*) = *ρ*(*x, t, TB*) is the density of cells with tumble bias *TB* at location *x* and time *t*, and *s* = *s*(*x, t*) is the concentration of attractant. The first equation describes how cell density changes due to chemotaxis and effective diffusion resulting from run-and-tumble behavior. *χ*(*TB*) and *μ*(*TB*) are the phenotype-dependent chemotactic (46) and diffusion coefficients characterizing cells’ chemotactic drift speed and random diffusive motion, respectively. *K*_*i*_ and *K*_*a*_ are the dissociation constants of the cells’ receptors for the attractant when in the inactive and active forms (47), which roughly set the lower and upper limits of concentration that the cells can detect. The second equation describes how the concentration of attractant *s* changes due to molecular diffusion *D*_*s*_ and cell consumption. *k* and *K*_*m*_ characterize consumption of the attractant by cells of all phenotypes. These PDEs are paired with boundary conditions stipulating that far ahead of the wave there are no cells and the concentration of attractant is *s*_∞_ (i.e. there is no attractant gradient ahead of the group): *ρ*(*x* → ∞, *t, TB*) = 0, ∂_*x*_*ρ*(*x* → ∞, *t, TB*) = 0, *s*(*x* → ∞, *t*) = *s*_∞_, and ∂_*x*_*s*(*x* → ∞, *t*) = 0.

We recently showed that cells of different phenotypes are able to migrate together by spatially organizing themselves in the group by chemotactic performance (12), which we quantify as *χ*(*TB*) because it determines how fast a phenotype climbs a static gradient (Fig. 1AB). Importantly, mounting evidence suggests that the mapping from phenotype *TB* to performance *χ* depends on the physical environment in which the cells swim. Experiments and simulations of chemotaxis in liquid have demonstrated that low-tumble bias cells have the highest chemotactic performance *χ* (24). On the other hand, experiments (4), theory (25), and simulations (48) indicate that cells with intermediate tumble bias (reorientation frequency) have the highest *χ*(*TB*) and *μ*(*TB*) in porous media.

**Figure 1:**
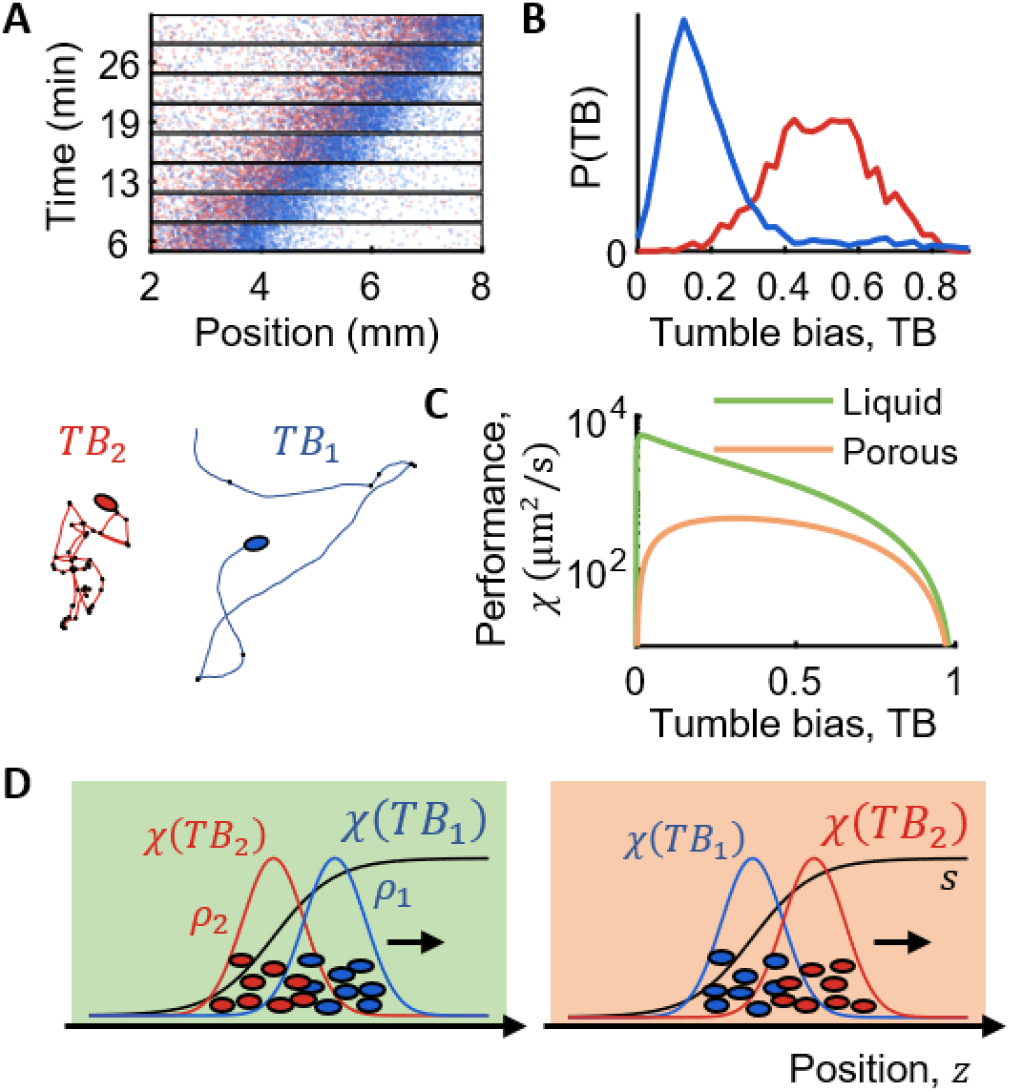
Chemotactic performance, and therefore spatial organization of phenotypes, is environment-dependent. **A,B)** Data from (12). Two isogenic populations with different phenotype distributions (quantified by tumble bias, TB) travel together in liquid by spontaneously sorting themselves within the migrating group. High-performing (low-TB, blue) cells are in front where the traveling gradient is shallow, and low-performing (high-TB, red) cells are in back where the traveling gradient is steep, equalizing chemotactic drift speed within the group. **C)** The chemotactic performance of each phenotype, quantified by χ, depends on the physical properties of the environment in which it swims. The highest-performing phenotype is different in liquid (green) and porous (orange) environments. **D)** Since cells in the group sort by performance, the spatial organization of phenotypes is different in different environments.

To explore the effects of a porous environment on group migration, we used a model for cell diffusion *μ*(*TB*) in porous media developed by Licata et al. (25) (Fig. 1C). In this model, runs that displace the cell by more than the mean free path length of the environment, *l*, are truncated to displacement *l*, and the cell idles in place until its next tumble. The effect of obstacles enters the diffusivity through one dimensionless parameter, *β*(*TB*) = *l λ*_*R*0_(*TB*)/*v*_0_ (SI Eqn. (4)). Here, *v*_0_ is the swimming speed during runs and *λ*_*R*0_(*TB*) is the phenotype-dependent average tumble rate, making *β*(*TB*) the ratio of the typical pore size and the average displacement of a run (25,48). Finally, for simplicity, we assumed that 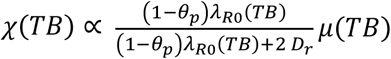, with environment-dependent proportionality constants that contain the details of chemotactic signal transduction and amplification. Here, the persistence of tumbles *θ*_*p*_ determines how much a tumble reorients the cell, and *D*_*r*_ is the rotational diffusion coefficient. The prefactor arises from the effects of rotational diffusion on chemotaxis (46,49–52), which has a substantial effect in liquid. This form is roughly justified because both *χ*(*TB*) and *μ*(*TB*) arise from cells’ run-and-tumble motility (14,15,53). This model predicts that cells with intermediate *TB* have the highest chemotactic coefficient *χ*(*TB*) in porous media because they tumble frequently enough to escape traps, while also maximizing the displacement of each run, consistent with recent agent-based simulations (48).

Since the mapping from phenotype to performance is environment-dependent, the spatial organization of phenotypes should also be environment-dependent. Numerical simulations of Eqns. (1)-(2) confirmed this prediction: in liquid, low tumble bias cells localize to the front of the group, and high tumble bias cells at the back; but in a porous environment, intermediate-tumble bias cells localize to the front, and both high- and low-tumble bias cells localize at the back (schematic in Fig. 1D; simulation results in Figs. S1-2). Thus, this model predicts that the spatial organization of swimming phenotypes in an isogenic population changes in different environments.

### Spatial organization exerts an environment-dependent selection pressure on swimming phenotype

What are the consequences of the spatial organization of phenotypes being environment-dependent? During migration, there is always a slow loss of cells behind the traveling group due to the cells’ finite sensitivity for the attractant that they chase (3,47,54,55). Simulations of Eqns. (1)-(2) revealed that by placing low-performers at the back of the group, spatial sorting causes them to fall behind the fastest. Furthermore, since the mapping from phenotype to performance depends on the environment, we find that different phenotypes fall behind fastest in liquid and porous environments. Over time, this differential leakage shifts the composition of the group towards phenotypes that are high-performing in each environment (Fig. 2A).

**Figure 2:**
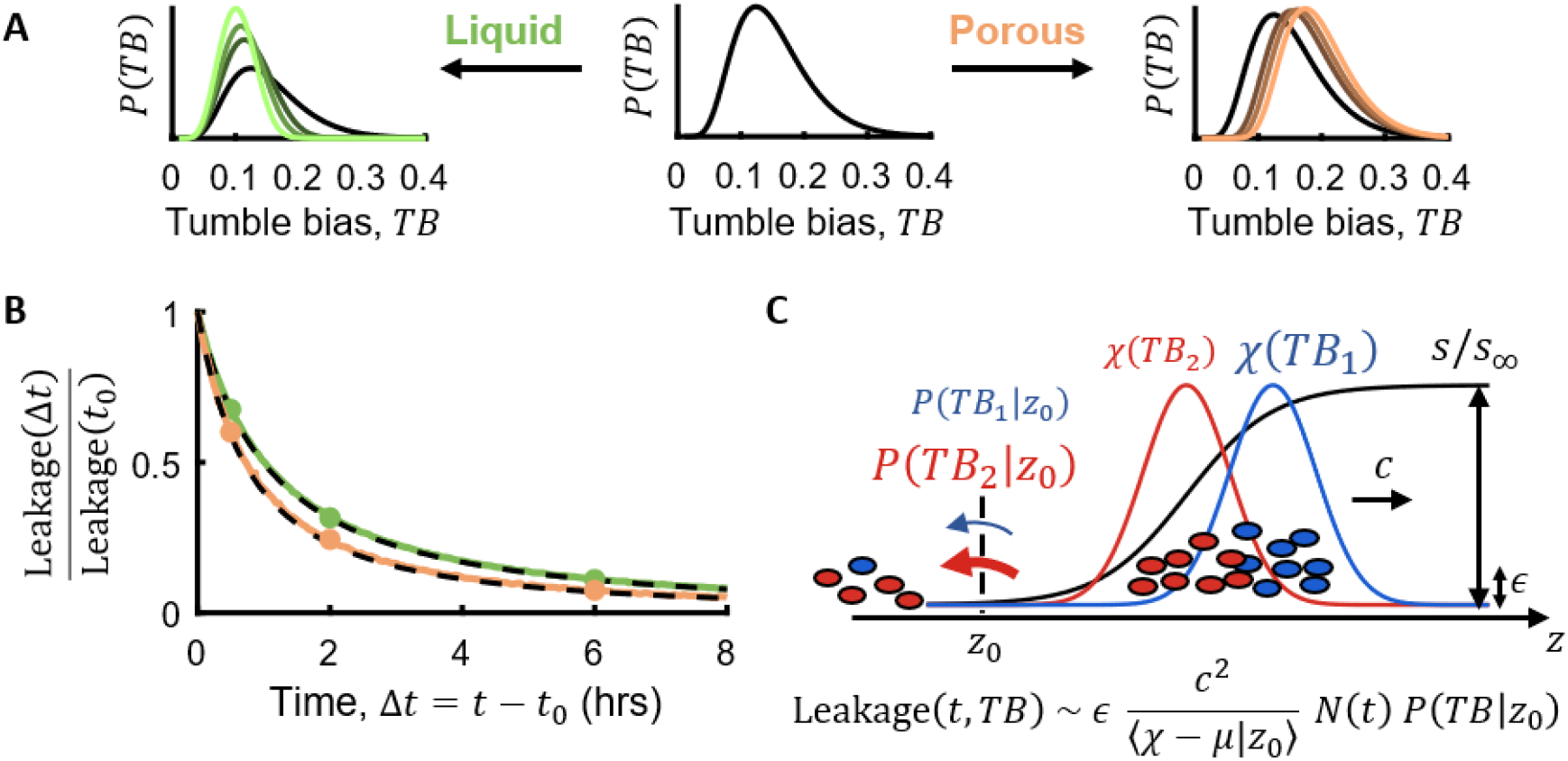
Spatial sorting causes low-performing phenotypes to fall behind the group. **A)** Since χ(TB) is environment-dependent, so is the differential loss of phenotypes. Thus, the migrating population creates its own selection pressure for high chemotactic performance. Black solid lines indicate the initial phenotype composition of two simulations, in liquid (green, left) and porous (orange, right) environments. Each line corresponds to a snapshot in time, marked by circles in (B), with lighter color corresponding to later time. The phenotype composition P(TB) of the same isogenic population shifts differently in liquid and porous environments. **B)** The theoretical expression for total leakage ∂_t_N(t) (Eqn. (6); black dashed lines) captures dependence of total leakage flux on time-dependent quantities (green: liquid; orange: porous; see Table S1 for parameter values). **C)** Schematic illustrating factors that contribute to total and differential loss of phenotypes off the back of the wave. Leakage depends on the cells’ sensitivity for the attractant ϵ = K_i_/s_∞_; the group migration speed c(t); the average chemotactic performance minus the average cell diffusivity at the back of the wave ⟨χ − μ|z_0_⟩; how localized the phenotype is at the back P(TB|z_0_); and the total number of cells traveling N(t), all of which except ϵ depend on time. Due to spatial sorting, low-performing cells (low χ(TB)) fall behind fastest because they are enriched at the back (high P(TB|z_0_)).

To better understand the factors affecting loss of cells from diverse migrating groups, we derived an approximate expression for the leakage flux of each phenotype. First, we transformed Eqns. (1)-(2) to a moving frame of reference 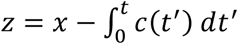, where *c*(*t*) is the group’s gradually slowing migration speed (SI). Then, with additional simplifications to make analytical progress, such as neglecting attractant diffusion (SI), we sought solutions for the density of each phenotype *ρ*(*z, t, TB*) and the attractant concentration *s*(*z, t*) that were approximately independent of time, finding:

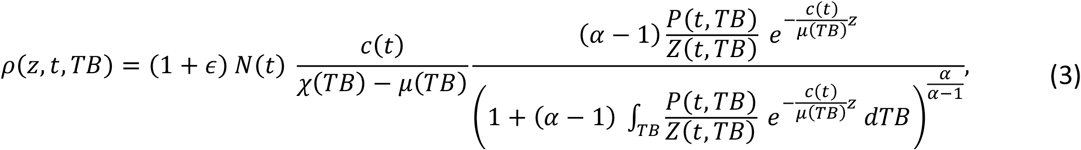

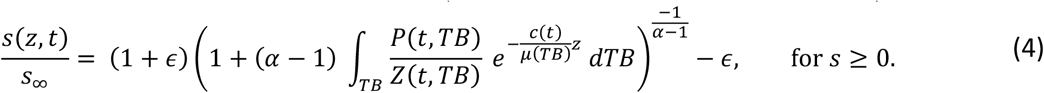

To arrive at these expressions, we had to assume 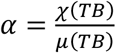 is constant for all phenotypes *TB*, but can be environment-dependent. *Z*(*t, TB*) is a factor that cannot be solved in closed form, but is fully determined by the composition of traveling phenotypes, *P*(*t, TB*) (SI). As in Keller and Segel’s classic result (2), the number of cells traveling (per cross-sectional area) sets the migration speed: *N*(*t*) = *c*(*t*) *s*_∞_/*k*. But unlike previous solutions (2,3), when cells have finite sensitivity for the attractant,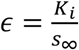, the attractant concentration *s* reaches zero at a finite position *z*_0_, which we define to be the back of the migrating group. We emphasize that Eqns. (3)-(4) are only approximations because they ignore changes in time, which are important at the back of the group.

Nevertheless, Eqns. (3)-(4) provide insights into the properties of diverse migrating groups. For example, since the traveling attractant gradient is generated by the cells’ own consumption and motion, its steepness is related to the phenotypes traveling. While the gradient steepness in traveling groups with a single phenotype is *λ* = *c*(*t*)/(*χ* − *μ*) (2,53), the phenotype composition in diverse groups varies with position *z*. Using Eqns. (3)-(4) we show that with diversity, *λ*(*z*) = *c*(*t*)/⟨*χ* − *μ*|*z*⟩ (SI), where ⟨*χ* − *μ*|*z*⟩ is the average of *χ* − *μ* over phenotypes located at position *z*. This result is important because it encapsulates the effect of phenotypic diversity on the shape of the traveling gradient: Towards the back of the group, the chemotactic abilities of the cells decrease and thus ⟨*χ* − *μ*|*z*⟩ gets smaller. These cells make a narrower and deeper dent in the attractant profile and a steeper local gradient. Thus, the spatial sorting mechanism (12) both generates and arises from local variations in the steepness of the traveling attractant gradient.

Next, to approximate the leakage flux, we noticed in simulations that there is a local minimum in cell density of each phenotype *ρ*(*z, t, TB*) that roughly coincides with *z*_0_, where the attractant concentration first equals zero behind the group. This means that cell diffusion negligible at *z*_0_. Furthermore, since there is no attractant at *z*_0_, physically there is no chemotactic flux there. Therefore, the leakage flux of each phenotype is ∂_*t*_*N*(*t, TB*) ∼ − *c*(*t*) *ρ*(*z*_0_, *t, TB*) = −*c*(*t*) *ρ*(*z*_0_, *t*) *P*(*TB*|*z*_0_, *t*) and the total leakage flux of all phenotypes is ∂_*t*_*N*(*t*) ∼ − *c*(*t*) *ρ*(*z*_0_, *t*). Although the quasi-steady state solutions Eqns. (3)-(4) do not allow for leakage, they do make a prediction for *ρ*(*z*_0_, *t*), from which we can determine the scale of the leakage flux. Doing this, we find:

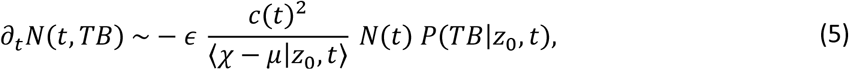

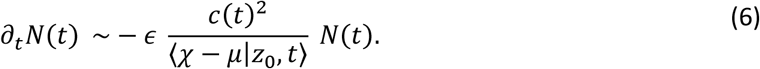

These expressions generalize previous results to account for multiple phenotypes traveling together (3,53). Due to spatial sorting, low performers contribute the most to ⟨*χ* − *μ*|*z*_0_, *t*⟩, which quantifies the average motility among cells located at *z*_0_, and thus the low performers set the scale of leakage for all phenotypes. The factor *P*(*TB*|*z*_0_, *t*) in Eqn. (5) indicates that lower-performing phenotypes, which localize more to the back of the group, fall behind faster. In the SI, use the quasi-steady state expression for *ρ*(*TB*), Eqn. (3), to show that low-performing phenotypes in the traveling group protect the high-performing ones from falling behind. The density of each traveling phenotype decays exponentially towards the back of the group. If a population with just a single, high-performing phenotype is migrating, its density *ρ*_*high*_(*TB*) decays with length scale *L*_*high*_ = (*χ* − *μ*)_*high*_/*c*(*t*). When a lower-performing phenotype with diffusivity is also traveling in the group, *L* _*high*_ is reduced by a factor of 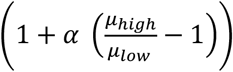. Thus, the presence of low performers in the group makes *ρ*_*high*_(*TB*) decay more rapidly, which exponentially reduces the density and leakage flux of the high performers at the back.

Due to this differential loss of phenotypes, the composition of the traveling group changes over time, slowly becoming enriched for phenotypes that are high-performing in the current environment. Thus, by sorting phenotypes in the group by their performance, collective migration creates a selective pressure for high performance in the environment that the population currently migrates through. However, leakage causes the group to slowly shrink and reduce speed, and therefore growth is needed to maintain migration (55). The long-term properties of the traveling group, and how it leverages diversity, depend on the balance between loss of cells due to leakage and production of cells due to growth.

### Minimal model of growth with non-genetic inheritance

To investigate the long-term properties of the migrating group, we constructed a minimal model of cell growth with multiple swimming phenotypes. Modeling growth with diversity requires specifying the extent to which daughter cells non-genetically inherit the phenotypes of their mothers (Fig. 3A). To make minimal assumptions, we modeled production of phenotypes as a Gaussian, auto-regressive process (SI), similar to recent work studying non-genetic inheritance of growth rate (56). Since tumble bias *TB* lies between 0 and 1, we defined *TB* = 1/(1 + *e*^*F*^) and instead modeled production of the transformed phenotype 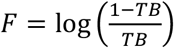. When the population is grown in batch culture, its distribution of phenotypes *F* approaches a Gaussian with population mean ⟨*F*⟩ and variance *σ*^2^. This model captures experimental measurements of the *TB* distribution in wild type *E. coli* cells (Fig. S3). Then, upon division, mother cells with phenotype *F*^′^ produce daughters cells with phenotypes *F* given by:

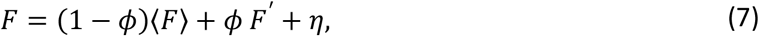

where *η* is a Gaussian random variable with mean ⟨*η*⟩ = 0 and variance ⟨*η*^2^⟩ = *σ*^2^ (1 − *ϕ*^2^), and *ϕ* is the correlation between mother and daughter phenotypes, *F*^′^ and *F*. When *ϕ* = 0, daughter cell phenotypes are chosen at random from the batch culture population. As *ϕ* → 1, the mean daughter phenotype becomes more similar to the mother’s phenotype, and the variation around the mean shrinks. Finally, we assumed that nutrients are abundant everywhere and distinct from the attractant, such that cells grow exponentially during migration (14,53). For simplicity, we assumed all that phenotypes divide with the same maximal rate *r* and we neglected changes in a cell’s swimming phenotype within its lifetime (26).

**Figure 3:**
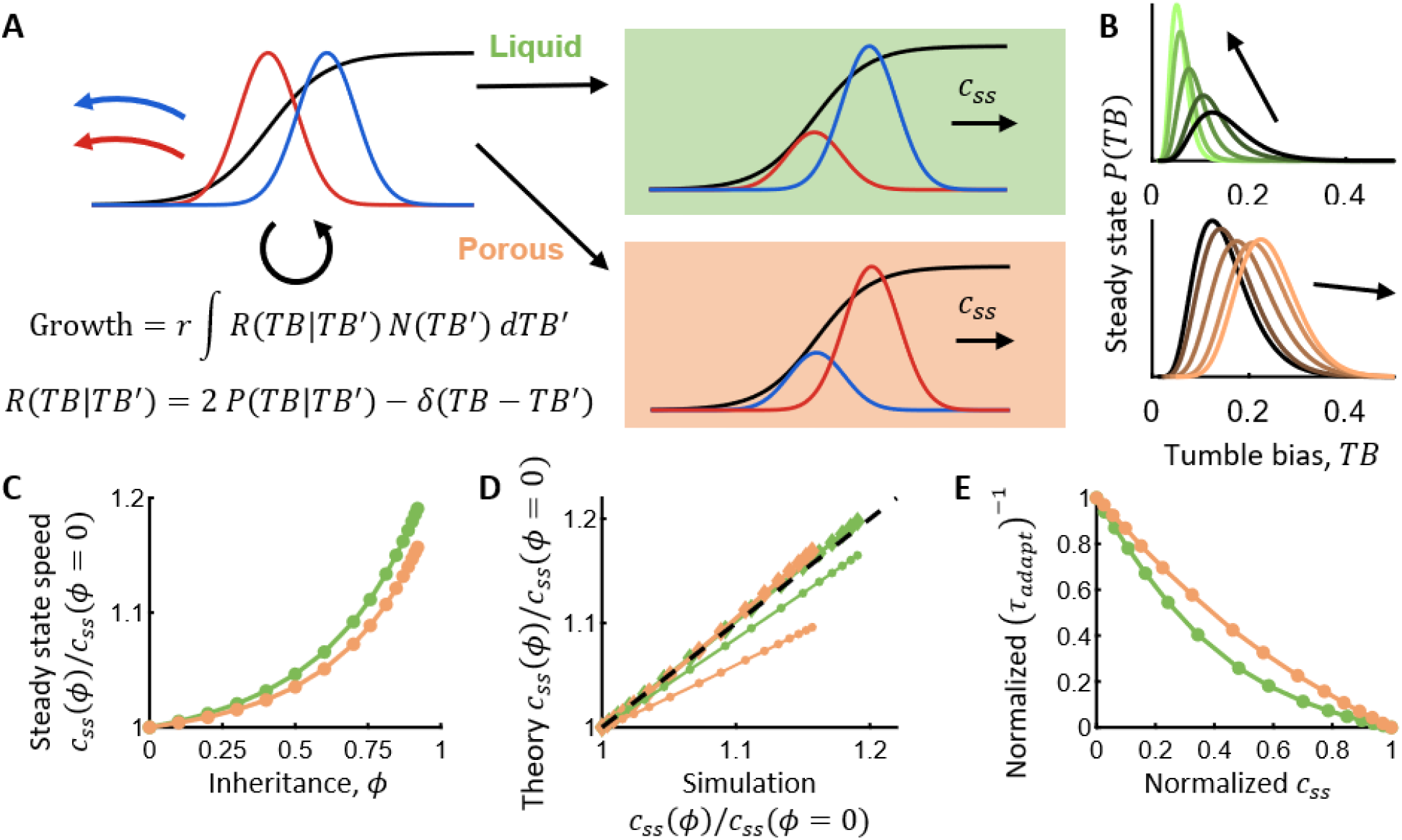
Growth and leakage balance to adapt the group’s composition to the environment, depending on non-genetic inheritance of phenotype. **A)** Growth is need to maintain migration over long distances by replacing lost cells. With multiple phenotypes, daughter cells phenotypes TB are drawn from a distribution P(TB|TB^′^) that depends on their mother’s phenotype TB^′^. This encodes the correlation between mother and daughter phenotypes, ϕ. Cells in the model divide at rate r, and divisions produce two daughters and remove the mother. With growth, the group composition P(TB) and migration speed c_ss_ reach steady states. Growth imperfectly replaces lost cells, causing the group composition to stably, dynamically shift towards high-performing phenotypes in each environment. **B)** Steady state phenotype compositions in groups migrating through liquid (top; green throughout) or porous (bottom; orange throughout) environments, for varying values of mother-daughter phenotype correlation ϕ, which increases from dark to light lines, or in the direction of the arrows. Black solid line: phenotype composition in batch culture, when not migrating. **C)** Larger shifts in composition caused by higher phenotypic inheritance lead to faster steady state migration speed, more cells traveling, and faster total population growth (14). **D)** Theory predicts that steady state migration speed is set by the lowest-performing phenotypes traveling, which depends on phenotypic inheritance, and it accurately captures steady state migration speeds in simulations. Each marker is a simulation of a population with different inheritance ϕ. Thick lines with diamonds plot the predictions of Eqn. (11), which accounts for attractant diffusion whose effect becomes noticeable in porous environments. Thin lines with circles are the predictions of Eqn. (10), which neglects attractant diffusion. Predictions in this panel took the steady state value of ⟨χ − μ|z_0_⟩ from simulations. **E)** Faster steady state migration trades off against time it takes the population to adapt to a new environment. Inverse adaptation time (τ_adapt_)_−1_ (or adaptation rate) and steady state migration speed c_ss_ from simulations were normalized to lie between 0 and 1. Larger values of ϕ lead to faster migration in both environments, but slower adaptation rate.

We can use this model to describe growth in the migrating group by adding a term on the right-hand side of Eqn. (1):

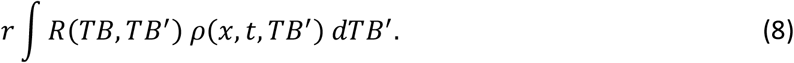

Here, *R*(*TB, TB*^′^) = 2 *P*(*TB*|*TB*^′^) − *δ*(*TB* − *TB*^′^) is a kernel that maps divisions of mother cells to production of daughters. *P*(*TB*|*TB*^′^) is the distribution of offspring phenotypes *TB* given that the mother’s phenotype *TB*^′^ that is implied by the dynamics in Eqn. (7) and by the mapping from *TB* to *F*. The factor of 2 accounts for production of two daughter cells upon division (whose phenotypes may be correlated with each other; SI), and the Dirac delta function accounts for removal of the mother cell from the population upon division. Using this model, we investigated how growth and leakage balance to determine the long-term properties of the migrating group.

### An isogenic population non-genetically adapts to migrate through multiple environments

Simulating our model of collective migration with growth, we found that the migrating group not only approaches a stable migration speed and population size, but also a stable phenotype composition (Fig. 3B). Leakage selectively removes low-performing phenotypes at the back of the group, but growth does not specifically replace the phenotypes that were lost. Therefore, the phenotype composition of the migrating group dynamically and stably shifted towards phenotypes that perform well in the current environment. This composition shift depended on the phenotypic inheritance parameter *ϕ*: with increasing *ϕ*, the composition shifted more because the traveling cells produce daughters that were more similar to themselves and more dissimilar to the cells that fell behind. In the limit of *ϕ* → 1, the migrating population would eventually purify to the single phenotype that performs best in the current environment.

These shifts in phenotype composition were accompanied by increases in steady state migration speed, population size, and total population growth (Fig. 3C; SI). Migration speed is set by the number of cells traveling, *c* ∝ *N*. Therefore, to understand why larger composition shifts lead to faster migration, we need to understand how the balance of growth and leakage determine group size at steady state.

Since growth is slow compared to migration (SI; (53)), we approximated the total leakage flux using Eqn. (6). If all phenotypes divide at roughly the same rate *r*, then the dynamics of the total number of cells traveling *N*(*t*) are (SI):

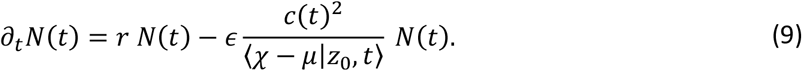

At steady state, growth and leakage balance, ∂_*t*_*N*(*t*) = 0, giving:

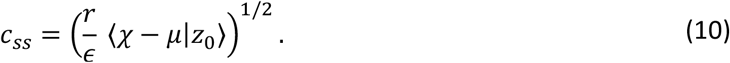

The expressions above neglected attractant diffusion, which is valid when the chemotactic abilities of the traveling cells are larger than the attractant diffusivity, *χ* − *μ* ≫ *D*_*s*_. While this is a good approximation in liquid, attractant diffusion cannot be neglected in porous environments.

To account for this, we also took an alternative approach based on recent theoretical work studying collective migration of a single phenotype (53), and extended it to include multiple phenotypes. In this approach, we made the ansatzes that, at the back of the group, the steady state attractant profile is *s*(*z*)/*s*_∞_ ∝ exp(∫ *λ*(*z*) *dz*), the total cell density profile is *ρ*(*z*) = *β*(*z*) (*ϵ* + *s*(*z*)/*s*_∞_), and the spatially-varying gradient steepness is *λ*(*z*) = *c*_*ss*_/⟨*χ* − *μ*|*z*_0_, *ss*⟩ (SI). With this approach, we arrive at the correction to Eqn. (10) with attractant diffusion:

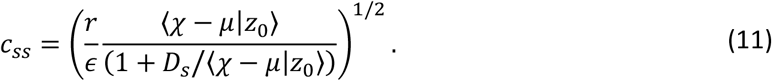

Surprisingly, the steady state migration speed of the entire group is set by the chemotactic abilities of cells at the back through ⟨*χ* − *μ*|*z*_0_⟩. With higher performers at the back, the steady state migration speed and population size increase. Therefore, higher inheritance *ϕ* and larger composition shifts lead to faster steady state migration by increasing the minimum performance and thus reducing the leakage rate (Fig. 3D). These effects could not be captured using the mean chemotactic abilities of the entire group, i.e. ⟨*χ* − *μ*⟩ (Fig. S4), highlighting the importance of the lowest performers traveling.

While higher inheritance drove faster steady state migration by causing larger shifts in phenotypic composition, it also slowed down the population’s response to a new environment. To quantify this trade-off, we extracted the relaxation time of migration speed near steady state for varying values of *ϕ* (Methods). Normalizing the adaptation rates and migration speeds to lie between 0 and 1 and plotting them against each other, we visualized the Pareto front of this trade-off (Fig 3E). The intuition for this trade-off can be understood by making an analogy to an Ornstein-Uhlenbeck (OU) process. In an OU process, a particle diffuses in an energy potential well, and the relaxation time of the particle’s position depends on its mass and the drag of the medium. A larger-mass particle with a shorter relaxation time is also less responsive to external forces. Likewise, lower inheritance shortens the relaxation time of the group’s phenotype composition but makes it less responsive to the selective pressure of collective migration.

Together, these results indicate that the selective pressure created by the population’s own collective migration, combined with cell growth, non-genetically adapts its phenotype composition to migrate effectively through multiple environments. Phenotypic inheritance modulates the extent of this non-genetic adaptation and thus the steady state wave speed, but trades off with slow responses to new environments. Importantly, fast migration provides an advantage to populations migrating through extended, nutrient-rich environments by more rapidly providing cells access to growth nutrients. Recent experiments and simulations have demonstrated that this leads to faster growth of the total population, both the cells in the migrating front and the cells left behind it (14) (SI). Thus, by adapting its phenotype composition to migrate faster, a diverse population increases its total growth.

### Non-genetic inheritance can enhance migration speed in varying environments

Why should a population maintain phenotypic diversity? A population that evolves to a single phenotype with the maximal value of *χ* − *μ* in a given environment would migrate fastest and thus grow its total population size fastest. But a migrating population can encounter multiple environments, where the highest-performing phenotype changes; this is when phenotypic diversity might be beneficial.

To quantify the role of diversity for navigating through multiple environments, we simulated diverse and non-diverse migrating populations that periodically encounter liquid and porous environments after fixed amounts of time *T* (Fig. 4). We fixed the time spent in each environment because traveling the same distance through a porous environment takes much longer than in liquid, and therefore the population would have much more time to adapt to the porous environment. For the population without diversity, we considered the generalist phenotype, defined as having the highest minimum value of *χ* − *μ* across the two environments (normalized by its maximum in each environment), and hence the highest minimum steady state migration speed (Fig. S5). Then, we generated diverse populations in which the most abundant phenotype was the generalist phenotype and the population variance *σ*^2^ was fixed, but the level of phenotypic inheritance *ϕ* was varied. Population performance was defined as the time-averaged migration speed across the two environments, after many environment switches.

**Figure 4:**
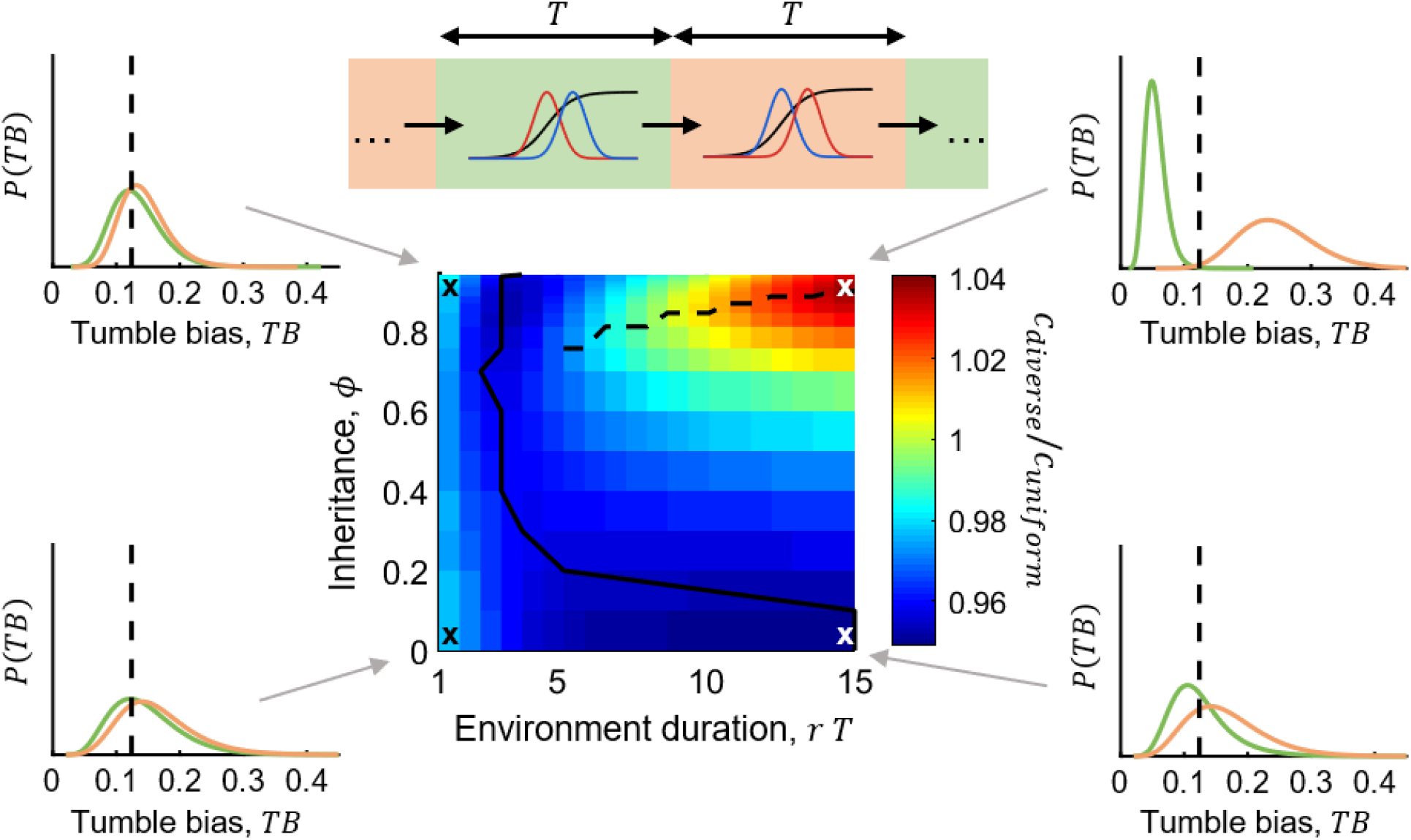
Diverse populations non-genetically adapt to migrate through varying environments, and can out-run a single-phenotype, generalist population in slowly-varying environments. Top: schematic of the simulation. Populations with varying mother-daughter phenotype correlation ϕ were simulated migrating through liquid and porous environments that switch after a fixed time T. Middle: after many environment switches, the stationary, time-averaged migration speed across the two environments c_diverse_ was computed and normalized by that of a single-phenotype, generalist population c_uniform_. Environment duration T is normalized by the cell division time 1/r. The diverse population adapts to varying environments in qualitatively different ways, depending on the time scale of environment changes, separated by the solid black line. Insets show the phenotype composition at the end of each liquid (green) and porous (orange) environment. Black dashed lines in the insets indicate the generalist phenotype. When the environment varies rapidly (r T < ∼3), and the population narrows its composition around the generalist phenotype, and larger ϕ increases migration speed by more effectively narrowing the phenotype composition. However, the diverse population never out-runs the generalist. When the environment varies slowly (r T > ∼3), the diverse population adapts its composition to each environment. An optimal amount of inheritance ϕ maximizes migration speed by balancing large composition shifts with the need to adapt before the environment switches. In this regime, a diverse population can out-run the generalist population.

The average migration speed of the diverse population exhibited two qualitatively different regimes (Fig. 4). In environments that varied quickly compared to the cells’ doubling time (*r T* < ∼3), higher inheritance *ϕ* monotonically increased the average migration speed. In this regime, the population composition narrowed around the generalist phenotype, and higher inheritance helped the population maintain the generalist phenotype (Fig. 4, left). However, the non-diverse population always migrated faster in these rapidly-changing environments. In slowly-varying environments (*r T* > ∼3), an optimal level of inheritance *ϕ* maximized the average migration speed (43). Here, the population composition tracked the environment, balancing the trade-off between achieving large shifts in composition with the need to adapt before the environment switched (Fig. 4, right). Once the period of each environment was sufficiently long, the diverse population out-performed the non-diverse one by shifting and narrowing its composition around the specialist phenotype in each environment. In general, the conditions in which the diverse population out-performs the generalist phenotype depend on the performance trade-offs between the particular environments that they encounter. Thus, the selective pressure imposed by differential leakage, paired with inheritance of phenotype, transiently and non-genetically adapts the population composition to migrate effectively through varying environments.

## Discussion

Populations of chemotactic bacteria can rapidly expand by chasing a gradient of attractant cue created by their own consumption (14,20), but the role of non-genetic diversity in this process has been unclear. Here, using analytical theory and simulations, we predict that diversity in swimming behaviors enables a migrating population to non-genetically adapt its phenotype composition to migrate through multiple physical environments. A key feature of this mechanism is the spatial organization of phenotypes during collective migration (12), which slowly removes phenotypes from the group that perform poorly in the current environment. To study this, we derived approximate analytical expressions for how the differential loss of phenotypes scales with group properties, and how these properties determine the steady state speed of diverse migrating populations. Furthermore, we found that the correlation between mother and daughter phenotypes controls a trade-off between strong adaption and slow responses to new environments. In slowly-varying environments, this adaptation mechanism enables a diverse population to out-run a non-diverse, generalist population. These results also demonstrate that collective behavior and phenotypic diversity can synergize to produce emergent functionalities (57).

The non-genetic adaptation mechanism we discovered here belongs to a broader class of bet-hedging strategies. A large body of work has studied diversification of growth rates among individuals (27–29,31– 34,37–40,42,45,58), and at least one study has considered the effect of diverse growth rates during undirected range expansion (44). In those cases, differences in growth rate select the fittest individuals in each environment, leading to faster population growth. But during directed range expansion considered here, faster population migration and growth require that the population select individuals with high chemotaxis performance, which doesn’t necessarily provide an intrinsic fitness advantage. We find that the population solves this problem by using the collective behavior of migration, itself, to create the selective pressure for high chemotaxis abilities.

In our model, cells with new phenotypes were produced by imperfect inheritance of their mother’s phenotype (56,43,30,59–62). While inheritance of phenotypes is mathematically identical to the well-studied process of stochastic phenotypic switching (32–34,63,36,41,64), it is conceptually and mechanistically different. Stochastic switching can require dedicated biochemical machinery, such as bistable switches (65,66), to produce a few discrete cell states. Non-genetic inheritance of a continuous phenotype, on the other hand, can readily arise in single-cell organisms, because variations in protein abundances and other molecules contribute to non-genetic differences, and daughter cells directly receive cytoplasmic contents from their mothers (67–69). There is evidence that phenotypic inheritance contributes to non-genetic adaptation of bacterial growth on formaldehyde (70), and non-genetic inheritance of swimming behavior has recently been measured in *E. coli* (26). Non-genetic adaptation may widely arise from imperfect inheritance of cellular components in single-celled organisms.

We investigated the effects of diversity in tumble bias, but bacteria also exhibit diversity in other phenotypic traits. These include, but are not limited to (71), swimming speed (23,52), receptor kinase adaptation time (52,72,73), receptor array composition (73–78), and attractant consumption rate. By influencing cells’ chemotaxis performance or the attractant gradient, these phenotypes can also become spatially sorted within the migrating group, and their organization can change in different physical and chemical environments. For example, recent work has measured spatial organization of Tar receptor expression in migrating groups chasing aspartate (21), and Tsr receptor (79) abundance is expected to be spatially sorted when the group chases serine. Furthermore, other physical environments, such as ones with different viscosities, can also give rise to different spatial organization of phenotypes (interestingly, *E. coli* were recently shown to maintain their tumble biases despite changes in environment viscosity (80)). Finally, for simplicity, we made various assumptions here about the relationship between *χ*(*TB*) and *μ*(*TB*). The cell-to-cell variations described above introduce additional variations in these quantities, even for cells with the same *TB*. However, they do not change our qualitative conclusions.

The non-genetic adaptation mechanism we predict here may play an important role during competition between multiple migrating genotypes (15,16). Non-genetic diversity could help a population with lower mean performance keep up with a higher-performing population by dynamically shifting its composition. Furthermore, low performers of the leading population may protect high performers of the trailing population. This could enable two genotypes to undergo stable co-migration, which has not yet been observed in experiments. Importantly, the approximate expressions for leakage derived here (Eqns. (5)-(6)) apply in these contexts, whether multiple strains of different genotypes travel together or multiple species travel together. This is because all the details of cell motility in Keller-Segel-like drift-diffusion models reduce to effective chemotactic and diffusion coefficients, and genotypes are only mathematically encoded by which individuals produce which upon division. More generally, these results emphasize that genotype-to-phenotype maps are not fixed, but dynamic and context-dependent. Furthermore, self-generated gradients drive collective migration of eukaryotic cells (81–83), and may even describe directed migration of higher organisms. Our results are expected to directly apply in these contexts, as well.

Beyond group migration, the conflicts between collective behavior and individuality (12,84–86), together with production of new individuals by growth, might generally enable populations to select individuals that are high-performing at the collective task, expanding our understanding of the ways populations can hedge their bets.

## Methods

Simulations were performed by converting the PDEs into a system of ODEs by discretizing space *x*. Spatial derivatives were computed using central differences. Integration forward in time was done using an explicit, 4^th^-order Runge-Kutta scheme (87,88). The initial condition was taken to be a steep sigmoidal function in space: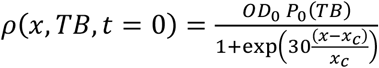, where *OD*_0_ was the initial density of cells, *P*_0_ (*TB*) is the initial composition, Θ(*x*) is the Heaviside step function, and *x*_*c*_ set the extent of the initial inoculation. We used 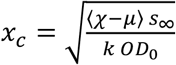, where the average in ⟨*χ* − *μ*⟩ is with respect to *P*_0_ (*TB*). This makes *x*_c_ roughly the characteristic length scale (width) of the wave that would be formed by the inoculated cells. For this choice of initial condition, nearly all phenotypes in the initial inoculation join the traveling group. *P*_0_(*TB*) was always the batch culture distribution. In simulations without growth, we used *OD*_0_ = 6; with growth in liquid, we used *OD*_0_ = 3; and with growth in porous media, we used *OD*_0_ = 0.5. The initial spatial grid size was *dx* = *x*_*c*_/50, using knowledge of the relevant length scale of the problem. In simulations of switching environments, the grid size was *dx* = *x*_*c*_/25. Time steps were *dt* = *dx*^2^/(4 *μ*_*max*_), where *μ*_*max*_ was the largest diffusion coefficient in the problem (including *D*_*s*_).

Simulations were done over a moving segment of space with no-flux boundary conditions on both sides. Several conditions were used to decide when to move the window of space being simulated. A new segment was simulated when: the attractant concentration at the right boundary dropped below a threshold; the peak cell density (and hence problem length scale) changed by more than 15%; the phenotype with the smallest or largest value of *TB* fell below a threshold (and could be removed from the simulation) or increased above a threshold (meaning that new phenotypes would imminently be produced by growth). If any of these conditions were met, the simulation was stopped, space far behind the wave peak was discarded, and the spatial region of the simulation was extended ahead of the direction of motion. The spatial grid size and time step were updated during this step, with 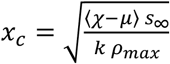, where the average was over phenotypes currently traveling, and *ρ*_*max*_ was the peak value of total cell density. The number of cells of each phenotype traveling was found by integrating over space from the location where the attractant *s* first reached zero behind the wave to space far ahead of the group. Then the cell density and attractant profiles were interpolated onto the new spatial grid, and the simulation was resumed.

To make predictions in Fig. 2B, we numerically computed *Z*(*t, TB*) in Eqn. (3) at each point in time, taking the migration speed *c*(*t*) and composition *P*(*t, TB*) as given from the simulation. *Z*(*t, TB*) was determined by requiring that the number of cells of each phenotype traveling *N*(*t, TB*) = *N*(*t*) *P*(*t, TB*) agreed with that in the simulation at time *t. N*(*t, TB*) in theory and simulations was computed by integrating cell density over the wave from *z* = *z*_0_ to *z* → ∞ (where 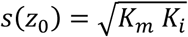 defines the back of the wave; see SI section **Steady state migration speed with attractant diffusion**). This determined *Z*(*t, TB*) up to a constant, which sets the location of *z* = 0 in the moving reference frame. To fix the constant, *Z*(*t, TB*) for the highest-performing phenotype was pinned at 1. We used Matlab’s *fmincon* function to find log(*Z*(*t, TB*)) numerically, since *Z*(*t, TB*) varied by orders of magnitude at any given time point. To help the optimization routine find solutions, we imposed a constraint on *Z*(*t, TB*) for consecutive values of *TB*: if *χ*(*TB*_*i*_) > *χ*(*TB*_*i*−1_), then we required that *Z*(*t, TB*_*i*_) < *Z*(*t, TB*_*i*−1_). Optimization was performed in two rounds for each time point. First, the sum of squared errors between *N*(*t, TB*) in the simulation and the analytical solution was minimized; then the sum of squared errors of log(*N*(*t, TB*)) was minimized to better capture *N*(*t, TB*) for low-abundance phenotypes, especially those at the back. For the first time point of the simulation, the optimization routine was initialized at *Z*(*t, TB*) = 1 for all *TB*, since this is the solution for one phenotype. In following time points, the optimization was initialized at the solution *Z*(*t, TB*) found for the previous time point, since *Z*(*t, TB*) is expected to change smoothly with time.

With cell growth, phenotypes that weren’t present at the start of the simulation could be produced. In this case, if the simulation of a spatial block stopped because the fraction of edge phenotypes increased above a threshold due to growth, then the range of phenotypes in the simulation was extended. Simulations were implemented with phenotype defined as *F*. The grid size of *F* was chosen to be 1/10^th^ of the standard deviation of the offspring phenotype distribution in order to resolve the growth matrix *R*(*F*|*F*^′^).

For the traveling populations to reach steady state in Fig. 3, group migration was simulated for 15 hours. The composition relaxation time was determined from the time course of migration speed as it approached steady state. Once the speed was within 10% of the steady state value, the speed profile was fit with an exponential function to extract the time scale of the slowest decaying mode, which was used to quantify the adaptation time.

Simulations in periodically varying environments were done by first simulating migration through one environment for the fixed environment duration. Then, to make minimal assumptions about how transitions between environments occurred, we took the cell density and attractant profiles at the end of the simulation in one environment and used those profiles as the initial conditions for a simulation in the next environment. The spatial grid size was adjusted for the expected change in speed in the new environment. In order for the simulation to reach a steady state, environments were alternated 15 times this way, for a total of 30 simulations for each parameter pair (*ϕ, T*). The time-averaged speed in each condition was computed by computing the total displacement over the last two environments (one liquid, one porous) and dividing by twice the duration of each environment, *T*.

All simulations were performed on Yale’s High-Performance Computing Clusters.

## Acknowledgements

We thank Jude Ong for collecting the tracking data in Fig. S3, and we thank Paul Turner and Sujit Datta for helpful discussions. This work was supported by NIH awards R01GM106189 (HM, TE), R01GM138533 (HM, TE), and F32GM131583 (HM).

## Contributions

HM and TE designed the research, performed theoretical analyses, and wrote the paper. HM wrote the code and ran the simulations.

