## Supplementary text and figures for "A mechanism for migrating bacterial populations to non-genetically adapt to new environments"

### Model of collective migration

We model collective migration without using an extension of the Keller-Segel model, similar to one developed recently (1). The main differences from Keller-Segel are: 1) we model co-migration of *E. coli* with different swimming phenotypes, quantified by tumble bias, $TB$; 2) cells' receptors have finite sensitivity for the chemoattractant, set by the receptor-ligand dissociation constant when the receptors are in the inactive state, $K_i$ (2).

The full PDEs for cell density $\rho(x,t,TB)$ and attractant concentration $s(x,t)$ are (reproducing Eqns. 1 and 2 of the main text):

$$\partial_t \rho(x,t,TB) = -\partial_x \left( \chi(TB)\, \rho(x,t,TB)\, \partial_x \log\left(\frac{1+\frac{s}{K_i}}{1+\frac{s}{K_a}}\right) - \mu(TB)\, \partial_x \rho(x,t,TB) \right), \quad (1)$$

$$\partial_t s(x,t) = D_s\, \partial_{xx} s(x,t) - k\, \frac{s}{K_m + s} \int_{TB} \rho(x,t,TB)\, dTB. \quad (2)$$

### Motility models in liquid and porous environments

Simulating collective migration requires models for $\chi(TB)$ and $\mu(TB)$ in each environment. Several previous works have shown that the diffusion coefficient $\mu(TB)$ resulting from run-and-tumble motion in 3D liquid is:

$$\mu_{liquid}(TB) = \frac{1}{3} \frac{v_0^2}{\left(1-\theta_p\right) \lambda_{R0}(TB) + 2\, D_r} (1 - TB), \quad (3)$$

where $v_0$ is the run speed, $\theta_p$ is the persistence of tumble (expectation of cosine of the angle between the cell's heading before and after a tumble) (3), $\lambda_{R0}(TB)$ is the average tumble rate in the absence of a gradient (inverse of the average run duration for exponentially distributed runs), and $D_r$ is the rotational diffusion coefficient. We assumed that $v_0$, $\theta$, and $D_r$ are independent of tumble bias $TB$ (4).

$\lambda_{R0}(TB)$ was described by a motor model developed previously (4–7), consisting of a free energy barrier of height $F$ between run and tumble states when the cell is adapted to the background concentration of attractant and a basal switching frequency $\omega$:

$$F(TB) = \log\left(\frac{(1-TB)}{TB}\right) \quad (4)$$

$$\lambda_{R0/T0}(TB) = \omega \exp\left(\mp \frac{1}{2} F(TB)\right), \quad (5)$$

where $\lambda_{T0}$ is the average tumble-to-run transition rate. Choosing a tumble bias phenotype $TB$ immediately fixes $F$, $\lambda_{R0}$, and $\lambda_{T0}$. It also fixes the baseline free energy difference $f_0$ between receptor active and inactive states and baseline receptor-associated CheA kinase activity $a_0$ in the absence of a gradient.

In all main text simulations, $\chi_{liquid}(TB) \propto \frac{(1-\theta_p)\lambda_{R0}(TB)}{(1-\theta_p)\lambda_{R0}(TB)+2\,D_r}\mu_{liquid}(TB)$, with the proportionality constant chosen such that $\alpha_{liquid} = \langle \chi_{liquid}(TB)/\mu_{liquid}(TB) \rangle = 10$ to match its value in a biophysical model, below. The expectation was taken with respect to the batch culture distribution of phenotypes (see section **Growth model**).

In porous environments, we used an expression for the diffusion coefficient derived by Licata et al. (8):

$$\mu_{porous}(TB) = \mu_{liquid}(TB)\left(1 - (1+\beta)\exp(-\beta) + \theta_p \exp(-\beta)\,(\beta + \exp(-\beta) - 1)\right). \quad (6)$$

Here, $\beta = l\,\lambda_{R0}/v_0$, where $l$ is the typical pore size of the porous mesh. To arrive at this expression, the authors assumed that runs that displaced the cell by more than $l$ were truncated at $l$, and the cell stalled in place until its next tumble. $l$ was chosen such that the population average diffusivity was about 50 $\mu m^2/s$, in line with recent measurements (9), giving $l \sim 30\ \mu m$. Finally, we assumed that $\chi_{agar}(TB) \propto \frac{(1-\theta_p)\lambda_{R0}(TB)}{(1-\theta_p)\lambda_{R0}(TB)+2\,D_r}\mu_{agar}(TB)$, with proportionality constant chosen such that $\alpha_{agar} = \langle \chi_{agar}(TB)/\mu_{agar}(TB) \rangle = 6.5$ (9,10).

To test whether the forms of $\chi(TB)$ we chose had qualitative effects on our results, we also modeled $\chi_{liquid}(TB)$ biophysically (4,6):

$$\chi_{liquid}(TB) = N\,\lambda'_R(f)_{f=f_0}\,\frac{\tau_m(TB)}{\left(1-\theta_p\right)\lambda_{R0}(TB) + 2\,D_r + \tau_m(TB)}\,\mu_{liquid}(TB). \quad (7)$$

Here, $N$ is the cooperativity of the receptors that sense the ligand, $\tau_m(TB)$ is the chemotaxis pathway adaptation time, and $\lambda'_R(f)_{f=f_0}$ is the gain of the tumble rate with respect to changes in receptor free energy $f$ in the adapted state $f_0$. The form of $\lambda'_R(f)_{f=f_0}$ and associated parameter values were taken from refs (6,7), and a phenomenological model for $\tau_m(TB)$ was taken from refs (1,11).

Simulations of migrating groups using this model showed the same qualitative behavior as described in the main text.

| Parameter | Value | Meaning | Source |
|---|---|---|---|
| $k$ | $0.126\ \mu\mathrm{M}\ \mathrm{OD}^{-1}\ s^{-1}$ | Attractant consumption rate | (1) |
| $D_s$ | $0$ or $800\ \mu\mathrm{m}^2/\mathrm{s}$ | Attractant diffusivity | (12,13) |
| $K_m$ | $0.5\ \mu\mathrm{M}$ | Monod constant | (1,14) |
| $s_\infty$ | $100\ \mu\mathrm{M}$ | Attractant concentration ahead of the wave | (1,9,15) |
| $K_i$ | $3.5\ \mu\mathrm{M}$ | Receptor lower sensing limit | This study, (1,2) |
| $K_a$ | $55\ \mathrm{mM}$ | Receptor upper sensing limit | (1) |
| $v_0$ | $25\ \mu\mathrm{m}/\mathrm{s}$ | Cell swimming speed | E.g. (4,16,17) |
| $\omega$ | $1.3\ \mathrm{s}^{-1}$ | Cell tumble rate parameter | (4,6,11,18) |
| $D_r$ | $0.062\ \mathrm{rad}^2/\mathrm{s}$ | Cell rotational diffusion coefficient | (19) |
| $\theta_p$ | 0.33 | Persistence of tumbles | (19) |
| $l$ | $30\ \mu\mathrm{m}$ | Average pore size | This study |
| $\alpha_{liquid}$ | 10 | $\langle \chi(TB)/\mu(TB) \rangle$ in liquid | This study |
| $\alpha_{porous}$ | 6.25 | $\langle \chi(TB)/\mu(TB) \rangle$ in porous media | (10) |
| $OD_0$ | 0.5, 3, or 6 OD | Initial cell density | This study |
| $\langle F \rangle$ | 1.9 | Batch culture mean phenotype | This study; Fig. S5 |
| $\sigma$ | 0.46 | Batch culture phenotype standard deviations | This study; Fig. S3 |
| $\phi$ | Varying | Mother-daughter correlation of phenotype $F$ | This study |
| $r$ | $0.83\ \mathrm{hr}^{-1}$ | Growth rate | E.g. (9) |

**Table S1: Parameter values used in simulations.**

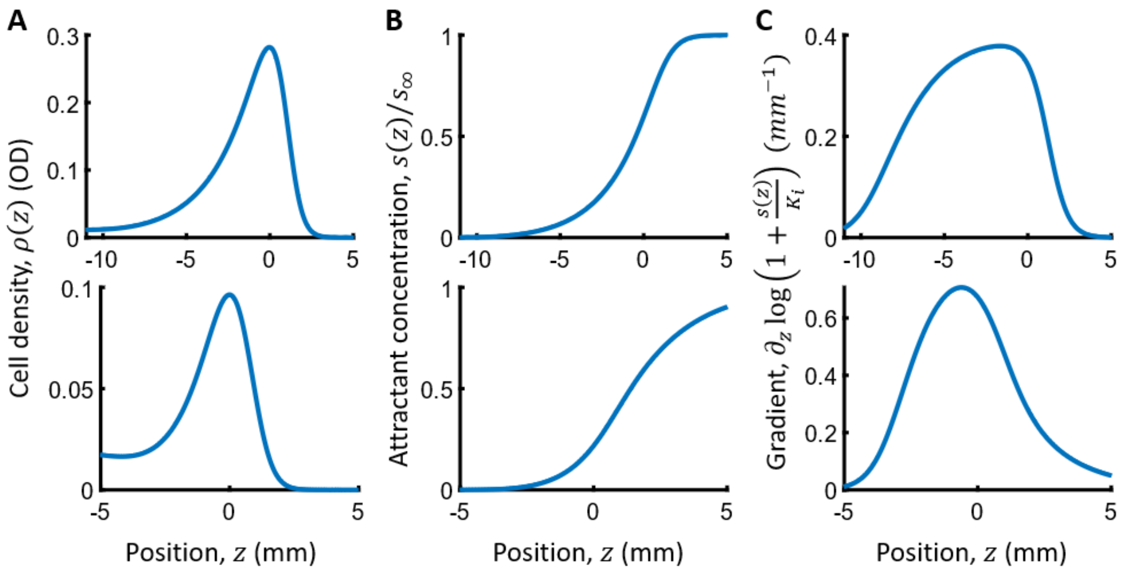

**Figure S1: Simulation profiles without growth.** Shown are simulation results for A) total cell density, B) attractant concentration, and C) perceived gradient steepness ($\partial_z \log(1 + s/K_i)$), in liquid (top) and porous (bottom) environments, at the last time point of simulations without growth. These are the same simulations used to generate the plots in Fig. 2 of the main text.

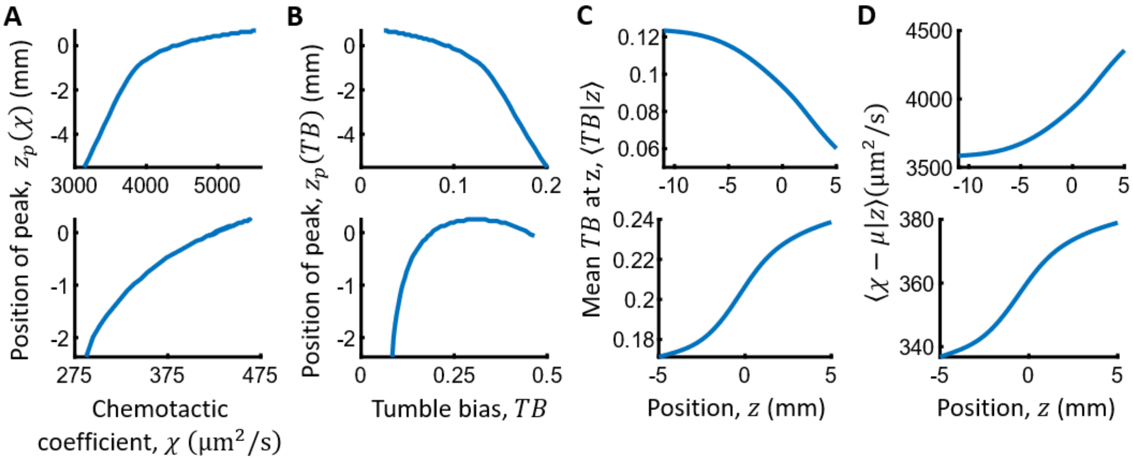

**Figure S2: Spatial organization is environment-dependent.** Shown are simulation results for the spatial organization of various quantities. A) Cells in the migrating group sort by chemotactic performance, regardless of environment (throughout, top: liquid, bottom: porous). Plotted is the position of the peak cell density of each phenotype $TB$ versus its chemotactic performance $\chi$. B) Since the mapping from phenotype to performance is environment-dependent, the organization of phenotypes is also environment-dependent. Shown is the peak cell density of each phenotype $TB$ versus $TB$, in each environment. C) Like B), the mean phenotype $TB$ at each position $z$ is sorted differently in liquid and agar. D) Like A), the mean motility $\langle\chi - \mu|z\rangle$ among cells at position $z$ is sorted the same (monotonically increasing with $z$) regardless of environment. These are the same simulations used to generate the plots in Fig. 2 of the main text.

### Quasi-steady state solutions

To simplify the PDEs above, we first assume that $K_m$ is small, justified by measurements putting its value below 1 μM (1,14). To make analytical progress, we also assume that $s \ll K_a$, the concentration of attractant is much smaller than the receptor-ligand dissociation constant when the receptors are in the active form. With these assumptions, the PDEs become:

$$\partial_t \rho(x,t,TB) = -\partial_x \left( \chi(TB)\, \partial_x \log\left(1 + \frac{s(x,t)}{K_i}\right) \rho(x,t,TB) - \mu(TB) \partial_x \rho(x,t,TB) \right) \quad (8)$$

$$\partial_t s(x,t) = D_s\, \partial_{xx} s(x,t) - k\, \Theta\big(s(x,t)\big) \int_{TB} \rho(x,t,TB)\, dTB, \quad (9)$$

where $\Theta(\dots)$ is the Heaviside step function, which ensures that attractant can only be consumed where it is present.

Next, we define a moving reference frame, $z = x - \int_0^t c(t')\, dt'$ and convert to the moving reference frame:

$$\begin{aligned} \partial_t \rho(z,t,TB) &- c(t)\, \partial_z \rho(z,t,TB) \\ &= -\partial_z \left( \chi(TB)\, \partial_z \log\left(1 + \frac{s(z,t)}{K_i}\right) \rho(z,t,TB) - \mu(TB) \partial_z \rho(z,t,TB) \right) \end{aligned} \quad (10)$$

$$\partial_t s(z,t) - c(t)\, \partial_z s(z,t) = D_s\, \partial_{zz} s(z,t) - k\, \Theta(s)\, \rho(z,t) \quad (11)$$

where $\rho(z,t) = \int_{TB} \rho(z,t,TB)\, dTB$.

For wave solutions, we seek a solution satisfying the following boundary conditions:

$$\rho(z,t,TB) \to 0 \text{ as } z \to \infty;\ \partial_z \rho(z,t,TB) \to 0 \text{ as } z \to \infty;\ s(z,t) \to s_\infty \text{ as } z \to \infty. \quad (12)$$

The equation for $\rho(z,t,TB)$ can also be written in terms of the fluxes $J(z,t,TB)$ of each phenotype in the moving reference frame:

$$\partial_t \rho(z,t,TB) = -\partial_z J(z,t,TB), \quad (13)$$

$$J(z,t,TB) = \left( \chi(TB)\, \partial_z \log\left(\epsilon + \frac{s(z,t)}{s_\infty}\right) - c(t) \right) \rho(z,t,TB) - \mu(TB)\, \partial_z \rho(z,t,TB), \quad (14)$$

where we have introduced $\epsilon = \frac{K_i}{s_\infty} \ll 1$.

Once the wave forms, it rapidly reaches a quasi-steady state; only on long time scales does the wave slow down due to loss of cells. Therefore, we can determine the quasi-steady wave solution by dropping the time derivatives (20).

First focusing on the equation for $\rho(z, t, TB)$, we integrate once with respect to $z$ and rearrange terms to get:

$$\partial_z \rho(z, t, TB) - \left( \frac{\chi(TB)}{\mu(TB)} \partial_z \log\left( \epsilon + \frac{s(z,t)}{s_\infty} \right) - \frac{c(t)}{\mu(TB)} \right) \rho(z, t, TB) = 0 \tag{15}$$

where, after applying the boundary conditions, the "constant" of integration is zero. This can be solved with an integrating factor

$$\exp\big(U(z, t, TB)\big) = \exp\left( \int_z - \left( \frac{\chi(TB)}{\mu(TB)} \partial_z \log\left( \epsilon + \frac{s(z,t)}{s_\infty} \right) - \frac{c(t)}{\mu(TB)} \right) dz \right) \tag{16}$$

$$= \exp\left( - \left( \frac{\chi(TB)}{\mu(TB)} \log\left( \epsilon + \frac{s(z,t)}{s_\infty} \right) - \frac{c(t)\, z}{\mu(TB)} \right) \right) \tag{17}$$

$U(z, t, TB)$ is analogous to an energy potential. Multiplying through by $\exp\big(U(z, t, TB)\big)$ and completing the total derivative gives:

$$\partial_z \Big( \exp\big(U(z, t, TB)\big)\ \rho(z, t, TB) \Big) = 0 \tag{18}$$

Integrating with respect to $z$ gives

$$\rho(z, t, TB) = Q(t, TB) \exp\big(-U(z, t, TB)\big) \tag{19}$$

where $Q(t, TB)$ comes from integration. The exact form of $Q(t, TB)$ cannot be solved in closed form, but is constrained implicitly by the distribution of phenotypes traveling at time $t$.

The equation for $s(z, t)$ is:

$$c(t)\ \partial_z s(z, t) = D_s\ \partial_{zz} s(z, t) - k\ \Theta(s)\ \rho(z, t). \tag{20}$$

Plugging in Eqn. (19) and dropping the Heaviside function, for now:

$$-c(t)\ \partial_z s(z, t) = D_s\ \partial_{zz} s(z, t) - k \int_{TB} Q(t, TB) \exp\big(-U(z, t, TB)\big)\ dTB \tag{21}$$

$$-c(t)\ \partial_z s(z, t) = D_s\ \partial_{zz} s(z, t) - k \int_{TB} Q(t, TB) \exp\left( \frac{\chi(TB)}{\mu(TB)} \log\left( \epsilon + \frac{s(z,t)}{s_\infty} \right) - \frac{c(t)\, z}{\mu(TB)} \right) dTB \tag{22}$$

Here, in order to proceed, we have to make the assumption that $\frac{\chi(TB)}{\mu(TB)} = \alpha$, a constant for all phenotypes. This assumption has been used before (9,10,21) and is based on the fact that directed and undirected motility both arise from the cells' run-and-tumble swimming behavior. But it ignores several kinds of cell-to-cell variations, such as variations in receptor array composition (22–27) and adaptation time (17,26,28). While this assumption is necessary to make analytical progress, as mentioned in this discussion, it does not affect our qualitative conclusions.

With this assumption, $s(z,t)$ can be separated from the integral over phenotypes:

$$-c(t)\,\partial_z s(z,t) = D_s\,\partial_{zz} s(z,t) - k\exp\left(\alpha\,\log\left(\epsilon + \frac{s(z,t)}{s_\infty}\right)\right)\int_{TB} Q(t,TB)\exp\left(-\frac{c(t)\,z}{\mu(TB)}\right)dTB \tag{23}$$

Rearranging:

$$\left(\epsilon + \frac{s(z,t)}{s_\infty}\right)^{-\alpha}\partial_z\left(\frac{s(z,t)}{s_\infty} + \frac{D_s}{c(t)}\partial_z\frac{s(z,t)}{s_\infty}\right) = \frac{k}{s_\infty\,c(t)}\int_{TB} Q(t,TB)\exp\left(-\frac{c(t)\,z}{\mu(TB)}\right)dTB \tag{24}$$

The terms inside the derivative on the left-hand side can be written

$$\left(\epsilon + \frac{s(z,t)}{s_\infty}\right)^{-\alpha}\partial_z\left(\frac{s(z,t)}{s_\infty}\left(1 + \frac{D_s}{c(t)}\partial_z\log\left(\frac{s(z,t)}{s_\infty}\right)\right)\right) = \frac{k}{s_\infty\,c(t)}\int_{TB} Q(t,TB)\exp\left(-\frac{c(t)\,z}{\mu(TB)}\right)dTB \tag{25}$$

This indicates that when $\frac{D_s}{c(t)} \ll \partial_z\log\left(\frac{s(z,t)}{s_\infty}\right)$, or the length scale of attractant diffusion in the wave is much smaller than the traveling gradient length scale, then attractant diffusion can be neglected. For now, we make this assumption.

Dropping the attractant diffusion term, and integrating with respect to $z$ gives:

$$\frac{1}{1-\alpha}\left(\epsilon + \frac{s(z,t)}{s_\infty}\right)^{1-\alpha} = -\frac{k}{s_\infty\,c(t)}\int_{TB}\frac{\mu(TB)}{c(t)}Q(t,TB)\exp\left(-\frac{c(t)\,z}{\mu(TB)}\right)dTB + A(t,TB) \tag{26}$$

Applying the boundary conditions, we find:

$$\frac{1}{1-\alpha}\,(\epsilon+1)^{1-\alpha} = A(t,TB). \tag{27}$$

Using this and rearranging Eqn. (26):

$$\left(\frac{\epsilon + \frac{s(z,t)}{s_\infty}}{1+\epsilon}\right)^{1-\alpha} = 1 + \frac{1}{(1+\epsilon)^{1-\alpha}}(\alpha-1)\frac{k}{s_\infty\,c(\tau t)}\int_{TB}\frac{\mu(TB)}{c(t)}Q(t,TB)\exp\left(-\frac{c(t)\,z}{\mu(TB)}\right)dTB. \tag{28}$$

Solving for $s(z,t)$:

$$\frac{s(z,t)}{s_\infty} = (1+\epsilon)\left(1+\frac{(\alpha-1)}{(1+\epsilon)^{1-\alpha}}\frac{k}{s_\infty\, c(t)}\int_{TB}\frac{\mu(TB)}{c(t)}Q(t,TB)\exp\left(-\frac{c(t)\,z}{\mu(TB)}\right)dTB\right)^{-\frac{1}{\alpha-1}} - \epsilon \qquad (29)$$

Finally, we redefine $Q(t,TB) \to \frac{c(t)}{\mu(TB)}\frac{s_\infty\, c(t)}{k}(1+\epsilon)^{1-\alpha}Q(t,TB)$ to get:

$$\frac{s(z,t)}{s_\infty} = (1+\epsilon)\left(1+(\alpha-1)\int_{TB}Q(t,TB)\exp\left(-\frac{c(t)\,z}{\mu(TB)}\right)dTB\right)^{-\frac{1}{\alpha-1}} - \epsilon \qquad (30)$$

Due to the Heaviside function in the approximate PDEs, this solution holds where $s(z,t) > 0$, i.e. for $z > z_0$; otherwise, $s(z,t) = 0$. Since we neglected attractant diffusion, this concentration profile has a kink at $z = z_0$, making its derivative discontinuous there. Eqn. (30) differs from the solution of Keller and Segel (29) and the zeroth-order solution of Novick-Cohen and Segel (20) in that $\epsilon$, the normalized sensitivity, is finite. This causes the attractant concentration to reach zero at a finite value of $z$.

As in Keller & Segel's original work, we can also integrate the equation for the attractant from $z = z_0$ to $z \to \infty$, giving:

$$c(t)\,(s_\infty - 0) = k\int_{z_0}^{\infty}\rho(z,t)\,dz \to c(t) = \frac{k}{s_\infty}N(t), \qquad (31)$$

where $N(t) = \int_{z_0}^{\infty}\rho(z,t)\,dz$ is the number of cells (per cross-sectional area) migrating at time $t$. As in Keller and Segel's model, the migration speed $c(t)$ depends on the number of cells traveling because migration is driven by the cells' consumption of the attractant.

Back to $\rho(z,t,TB)$, in the region where $s(z,t) > 0$ and $z > z_0$, we have, with the new definition of $Q(t,TB)$:

$$\rho(z,t,TB) = \frac{c(t)}{\mu(TB)}\frac{s_\infty\, c(t)}{k}(1+\epsilon)^{1-\alpha}Q(t,TB)\exp\big(-U(z,t,TB)\big) \qquad (32)$$

$$= \frac{c(t)}{\mu(TB)}\frac{s_\infty\, c(t)}{k}(1+\epsilon)^{1-\alpha}\,Q(t,TB)\,\left(\epsilon+\frac{s_0(z,t)}{s_\infty}\right)^{\alpha}\exp\left(-\frac{c(t)\,z}{\mu(TB)}\right) \qquad (33)$$

$$= (1+\epsilon)\frac{c(t)}{\chi(TB)-\mu(TB)}N(t)\frac{(\alpha-1)\,Q(t,TB)\exp\left(-\frac{c(t)\,z}{\mu(TB)}\right)}{\left(1+(\alpha-1)\,\int_{TB}Q(t,TB)\exp\left(-\frac{c(t)\,z}{\mu(TB)}\right)dTB\right)^{\frac{\alpha}{\alpha-1}}}. \qquad (34)$$

We chose not lump $(\alpha-1)$ into the definition of $Q(t,TB)$. Keeping them separate tends to make the peak cell density in the group roughly coincide with $z = 0$, separating a diffusive region in the front of the group where there is almost no gradient ($z > 0$) and a chemotactic region in the middle/back of the group ($z < 0$). The solution above for $\rho(z,\tau,TB)$ is only defined for $z > z_0$. At $z_0$, where the attractant $s = 0$, cell density is finite. We will use this later to make predictions about the leakage flux at the back of the wave. The dimensions of $\rho(z,t,TB)$ come from the prefactor, $\frac{c(\tau)}{\chi(TB)-\mu(TB)}N(t)$. $\frac{\chi(TB)-\mu(TB)}{c(t)}$ is roughly the width of the density profile of phenotype $TB$.

The distribution of phenotypes traveling is contained in $Q(t,TB)$, which is constrained by $\int_{z_0}^{\infty}\rho(z,t,TB)\,dz = N(t,TB)$, where $N(t,TB)$ is the number of cells of each phenotype traveling at time $t$. Without loss of generality, we can write $Q(t,TB) = P(\tau,TB)/Z(t,TB)$.

$$\rho(z,t,TB) = (1+\epsilon)\frac{c(t)}{\chi(TB)-\mu(TB)}N(t)\frac{(\alpha-1)\frac{P(t,TB)}{Z(t,TB)}\exp\left(-\frac{c(t)\,z}{\mu(TB)}\right)}{\left(1+(\alpha-1)\int_{TB}\frac{P(t,TB)}{Z(t,TB)}\exp\left(-\frac{c(t)\,z}{\mu(TB)}\right)dTB\right)^{\frac{\alpha}{\alpha-1}}} \quad (35)$$

From $\int_{z_0}^{\infty}\rho(z,t,TB)\,dz = N(t,TB)$, and for a given traveling distribution $P(t,TB) = N(t,TB)/N(t)$, the quantity $Z(t,TB)$ (and hence $Q(t,TB)$) is fixed by the implicit solution to:

$$\int_{z_0}^{\infty}(1+\epsilon)\frac{c(t)}{\chi(TB)-\mu(TB)}N(t)\frac{(\alpha-1)\frac{P(t,TB)}{Z(t,TB)}\exp\left(-\frac{c(t)\,z}{\mu(TB)}\right)}{\left(1+(\alpha-1)\int_{TB}\frac{P(t,TB)}{Z(t,TB)}\exp\left(-\frac{c(t)\,z}{\mu(TB)}\right)dTB\right)^{\frac{\alpha}{\alpha-1}}}\,dz = N(t,TB). \quad (36)$$

### Traveling gradient steepness

We must be clear that these solutions, Eqns. (30) and (35), have serious issues. Most importantly, the wave is not at steady state, and time derivatives at the back cannot be ignored. As a result, cell density cannot be described by an energy potential (Eqn. (19)). Still, they are useful for understanding the properties and organization of the wave, particularly in the region where chemotaxis dominates, roughly corresponding to $z < 0$. We use them here to derive the steepness of the traveling gradient the cells chase.

First, the density of each phenotype can be written as:

$$\rho(z,t,TB) = \frac{N(t)\,c(t)}{\chi(TB)-\mu(TB)}\frac{(\alpha-1)\frac{P(t,TB)}{Z(t,TB)}\exp\left(-\frac{c(t)\,z}{\mu(TB)}\right)}{\left(1+(\alpha-1)\int_{TB}\frac{P(t,TB)}{Z(t,TB)}\exp\left(-\frac{c(t)\,z}{\mu(TB)}\right)dTB\right)}\left(\epsilon+\frac{s(z,t)}{s_\infty}\right). \quad (37)$$

Towards the back of the group where chemotaxis dominates, the exponential term in Eqn. (35) is much larger than 1, giving:

$$\rho(z,t,TB) \sim \frac{c(t)}{\chi(TB)-\mu(TB)}N(t)\frac{\frac{P(t,TB)}{Z(t,TB)}\exp\left(-\frac{c(t)\,z}{\mu(TB)}\right)}{\int_{TB}\frac{P(t,TB)}{Z(t,TB)}\exp\left(-\frac{c(t)\,z}{\mu(TB)}\right)dTB}\left(\epsilon+\frac{s(z,t)}{s_\infty}\right). \quad (38)$$

Next, the total cell density profile in the wave is

$$\rho(z,t) = \int_{TB} \rho(z,t,TB)\, dTB \tag{39}$$

$$\sim\ c(t)\, N(t) \frac{\int_{TB} \frac{1}{\chi(TB)-\mu(TB)} \frac{P(t,TB)}{Z(t,TB)} \exp\left(-\frac{c(t)\, z}{\mu(TB)}\right) dTB}{\int_{TB} \frac{P(t,TB)}{Z(t,TB)} \exp\left(-\frac{c(t)\, z}{\mu(TB)}\right) dTB} \left(\epsilon + \frac{s(z,t)}{s_\infty}\right). \tag{40}$$

Using Eqns. (38) and (40), the local composition of phenotypes at position $z$, $P(TB|z,t)$, is:

$$P(TB|z,t) = \frac{\rho(z,t,TB)}{\rho(z,t)} = \frac{\frac{1}{\chi(TB)-\mu(TB)} \frac{P(t,TB)}{Z(t,TB)} \exp\left(-\frac{c(t)\, z}{\mu(TB)}\right)}{\int_{TB} \frac{1}{\chi(TB)-\mu(TB)} \frac{P(t,TB)}{Z(t,TB)} \exp\left(-\frac{c(t)\, z}{\mu(TB)}\right) dTB}, \tag{41}$$

and the mean value of $\chi(TB) - \mu(TB)$ among cells at position $z$ is:

$$\langle \chi - \mu | z,t \rangle = \int_{TB} \big(\chi(TB) - \mu(TB)\big) P(TB|z,t) dTB \tag{42}$$

$$\sim \frac{\int_{TB} \frac{P(t,TB)}{Z(t,TB)} \exp\left(-\frac{c(t)\, z}{\mu(TB)}\right) dTB}{\int_{TB} \frac{1}{\chi(TB)-\mu(TB)} \frac{P(t,TB)}{Z(t,TB)} \exp\left(-\frac{c(t)\, z}{\mu(TB)}\right) dTB}. \tag{43}$$

This lets us write the equations for cell density at the back of the wave in simpler forms:

$$\rho(z,t,TB) \sim\ N(t) \frac{c(t)}{\langle \chi - \mu | z,t \rangle} P(TB|z,t) \left(\epsilon + \frac{s(z,t)}{s_\infty}\right) \tag{44}$$

and

$$\rho(z,t) \sim N(t) \frac{c(t)}{\langle \chi - \mu | z,t \rangle} \left(\epsilon + \frac{s(z,t)}{s_\infty}\right). \tag{45}$$

From this, we can readily read off that $L(z,t) = \langle \chi - \mu | z,t \rangle / c(t)$ is the typical length scale of the group at position $z$.

Furthermore, this form of the quasi-steady state solution makes it straightforward to determine the traveling gradient steepness. We will find that the local phenotype composition at position $z$ sets the local gradient steepness.

$$\lambda(z,t) = \partial_z \log\left(\epsilon + \frac{s(z,t)}{s_\infty}\right) = \frac{1}{\left(\epsilon + \frac{s(z,t)}{s_\infty}\right)} \partial_z \left(\frac{s(z,t)}{s_\infty}\right) \tag{46}$$

From Eqn. (20) (with $D_S \to 0$), we have $\partial_z \left(\frac{s(z,t)}{s_\infty}\right) = \frac{1}{N(t)} \rho(z,t)$. Together with Eqn. (45), this means the local gradient steepness $\lambda(z,t)$ towards the back where chemotaxis dominates is:

$$\lambda(z,t) = \frac{c(t)}{\langle \chi - \mu | z,t \rangle}, \tag{47}$$

which is exactly the inverse length scale of the migrating group, $L(z,t)$. The gradient length scale is the distance over which concentration drops, which is the width of the wave.

To understand this, remember that the gradient is created by the cells' own consumption. Since all cells are assumed to consume the attractant at the same rate, a cell with low diffusivity creates a "deep" and "narrow" dent in the attractant concentration profile, while a high diffusivity cell creates a "shallow" and "wide" dent. This is reflected above, where the perceived gradient steepness at each location is inversely proportional to $\langle \chi - \mu | z,t \rangle$. Cells at the back have smaller $\chi(TB)$ and $\mu(TB)$; therefore, they make the gradient steeper there. Additionally, $\partial_z \log\left(\epsilon + \frac{s(z,t)}{s_\infty}\right)$ scales with the wave speed $c(t)$ – with a fixed phenotype composition, migrating faster requires that gradient is steeper.

### Low performers exponentially reduce the density of high performers at the back of the group

We can also use the quasi-steady state solutions Eqns. (30) and (35) to get a sense for the relative abundances of different phenotypes at the back of the group. Say there are two phenotypes traveling together, $TB_1$ and $TB_2$. The density of $TB_1$ is:

$$\rho(z,t,TB_1) = \frac{(1+\epsilon)\,N(t)\,c(t)}{\chi(TB) - \mu(TB)} \frac{(\alpha-1)\left(\frac{P(t,TB)}{Z(t,TB)}\exp\left(-\frac{c(t)\,z}{\mu(TB)}\right)\right)_1}{\left(1 + (\alpha-1)\left(\frac{P(t,TB)}{Z(t,TB)}\exp\left(-\frac{c(t)\,z}{\mu(TB)}\right)\right)_1 + (\alpha-1)\left(\frac{P(t,TB)}{Z(t,TB)}\exp\left(-\frac{c(t)\,z}{\mu(TB)}\right)\right)_2\right)^{\frac{\alpha}{\alpha-1}}}. \tag{48}$$

Towards the back of the wave, the exponential terms in the denominator dominate, so we can drop the 1:

$$\rho(z,t,TB_1) \sim \frac{(1+\epsilon)\,N(t)\,c(t)}{(\chi(TB) - \mu(TB))(\alpha-1)^{\frac{1}{\alpha-1}}} \frac{\left(\frac{P(t,TB)}{Z(t,TB)}\exp\left(-\frac{c(t)\,z}{\mu(TB)}\right)\right)_1}{\left(\left(\frac{P(t,TB)}{Z(t,TB)}\exp\left(-\frac{c(t)\,z}{\mu(TB)}\right)\right)_1 + \left(\frac{P(t,TB)}{Z(t,TB)}\exp\left(-\frac{c(t)\,z}{\mu(TB)}\right)\right)_2\right)^{\frac{\alpha}{\alpha-1}}}. \tag{49}$$

We can write this as:

$$\rho(z,t,TB_1) = \frac{(1+\epsilon)\,N(t)\,c(t)}{(\chi(TB) - \mu(TB))(\alpha-1)^{\frac{1}{\alpha-1}}} \frac{\left(\frac{P(t,TB)}{Z(t,TB)}\exp\left(-\frac{c(t)\,z}{\mu(TB)}\right)\right)_1 / \left(\frac{P(t,TB)}{Z(t,TB)}\exp\left(-\frac{c(t)\,z}{\mu(TB)}\right)\right)_1^{\frac{\alpha}{\alpha-1}}}{\left(1 + \left(\frac{P(t,TB)}{Z(t,TB)}\exp\left(-\frac{c(t)\,z}{\mu(TB)}\right)\right)_2 / \left(\frac{P(t,TB)}{Z(t,TB)}\exp\left(-\frac{c(t)\,z}{\mu(TB)}\right)\right)_1\right)^{\frac{\alpha}{\alpha-1}}} \tag{50}$$

or:

$$\rho(z,t,TB_1) = \frac{(1+\epsilon)\,N(t)\,c(t)}{\big(\chi(TB)-\mu(TB)\big)(\alpha-1)^{\frac{1}{\alpha-1}}} \frac{\left(\left(\frac{P(t,TB)}{Z(t,TB)}\right)^{-\frac{1}{\alpha-1}} \exp\left(\frac{c(t)\,z}{(\alpha-1)\mu(TB)}\right)\right)_1}{\left(1+\frac{\left(\frac{P(t,TB)}{Z(t,TB)}\right)_1}{\left(\frac{P(t,TB)}{Z(t,TB)}\right)_2} \exp\left(-\left(\frac{c(t)}{\mu(TB_2)}-\frac{c(t)}{\mu(TB_1)}\right)z\right)\right)^{\frac{\alpha}{\alpha-1}}}. \tag{51}$$

Then, say $\chi(TB_1) < \chi(TB_2)$ and $\mu(TB_1) < \mu(TB_2)$. Since $\frac{1}{\mu(TB_1)} > \frac{1}{\mu(TB_2)}$, towards the back the exponential in the denominator goes to zero, leaving:

$$\rho(z,t,TB_1) \sim \frac{(1+\epsilon)\,N(t)\,c(t)}{\big(\chi(TB)-\mu(TB)\big)(\alpha-1)^{\frac{1}{\alpha-1}}} \left(\left(\frac{P(t,TB)}{Z(t,TB)}\right)^{-\frac{1}{\alpha-1}} \exp\left(\frac{c(t)\,z}{(\alpha-1)\mu(TB)}\right)\right)_1. \tag{52}$$

This is the behavior of cell density if there is just one phenotype traveling: towards the back, cell density decays with length scale

$$L(TB_1) = \frac{(\alpha-1)\,\mu}{c} = \frac{(\chi-\mu)}{c}. \tag{53}$$

Then, the density of the high-performing phenotype is:

$$\rho(z,t,TB_2) = \frac{(1+\epsilon)\,N(t)\,c(t)}{\big(\chi(TB)-\mu(TB)\big)(\alpha-1)^{\frac{1}{\alpha-1}}} \frac{\left(\left(\frac{P(t,TB)}{Z(t,TB)}\right)^{-\frac{1}{\alpha-1}} \exp\left(\frac{c(t)\,z}{(\alpha-1)\mu(TB)}\right)\right)_2}{\left(1+\frac{\left(\frac{P(t,TB)}{Z(t,TB)}\right)_2}{\left(\frac{P(t,TB)}{Z(t,TB)}\right)_1} \exp\left(-\left(\frac{c(t)}{\mu(TB_1)}-\frac{c(t)}{\mu(TB_2)}\right)z\right)\right)^{\frac{\alpha}{\alpha-1}}}. \tag{54}$$

Towards the back, the exponential in the denominator gets large, giving:

$$\rho(z,t,TB_2) \sim \frac{(1+\epsilon)\,N(t)\,c(t)}{\big(\chi(TB)-\mu(TB)\big)(\alpha-1)^{\frac{1}{\alpha-1}}} \frac{\left(\left(\frac{P(t,TB)}{Z(t,TB)}\right)^{-\frac{1}{\alpha-1}} \exp\left(\frac{c(t)\,z}{(\alpha-1)\mu(TB)}\right)\right)_2}{\left(\frac{\left(\frac{P(t,TB)}{Z(t,TB)}\right)_2}{\left(\frac{P(t,TB)}{Z(t,TB)}\right)_1} \exp\left(-\left(\frac{c(t)}{\mu(TB_1)}-\frac{c(t)}{\mu(TB_2)}\right)z\right)\right)^{\frac{\alpha}{\alpha-1}}}. \tag{55}$$

which simplifies to:

$$\rho(z,t,TB_2) = \frac{(1+\epsilon)\,N(t)\,c(t)\,\left(\frac{P(t,TB)}{Z(t,TB)}\right)_2 \left(\frac{P(t,TB)}{Z(t,TB)}\right)_1^{\frac{\alpha}{\alpha-1}}}{\big(\chi(TB)-\mu(TB)\big)(\alpha-1)^{\frac{1}{\alpha-1}}} \exp\left(\left(\frac{\alpha}{\alpha-1}\frac{c(t)}{\mu(TB_1)}-\frac{c(t)}{\mu(TB_2)}\right)z\right). \tag{56}$$

Since $\frac{1}{\mu(TB_1)} > \frac{1}{\mu(TB_2)}$ and $\frac{\alpha}{\alpha-1} > 1$, this means that $\left(\frac{\alpha}{\alpha-1}\frac{c(t)}{\mu(TB_1)} - \frac{c(t)}{\mu(TB_2)}\right) > 0$, and the density of the high-performing phenotype decays towards the back (as $z \to -\infty$). The length scale of this decay is:

$$L(TB_2) = \frac{1}{\left(\frac{\alpha}{\alpha-1}\frac{c(t)}{\mu(TB_1)} - \frac{c(t)}{\mu(TB_2)}\right)} = \frac{(\alpha-1)\mu(TB_2)}{c(t)\left(1+\alpha\left(\frac{\mu(TB_2)}{\mu(TB_1)}-1\right)\right)} < \frac{(\alpha-1)\,\mu(TB_2)}{c(t)} \tag{57}$$

where the last step follows because $\mu(TB_2) > \mu(TB_1)$ and $\alpha > 1$. This means that density profile of the high performing phenotype decays much faster when a lower performer is present, which exponentially reduces its density at the back.

### Leakage flux

To derive an approximate leakage flux, we must break our assumption that the wave is at (quasi) steady state. By integrating the PDE for cell density from $z = z_0$ to $z \to \infty$, keeping the time derivative term, we get that the rate of change of the number of cells in the wave is:

$$\partial_t N(t,TB) = \int_{z0}^{\infty} \partial_t \rho(z,t,TB)\,dz = J(z_0,t,TB) \tag{58}$$

At $z_0$, there is no attractant, so physically the chemotactic flux should be zero. This is the first break from the quasi-steady solution, which requires that there be chemotactic flux at $z_0$ in order for there to be no changes in time. Furthermore, in simulations we noticed that at $z_0$ there is a local minimum in cell density: behind $z_0$ are cells that were leaked at an earlier time, when the leakage flux was higher; ahead of $z_0$ is the increasing cell density of the wave. Therefore, we also neglect the diffusive flux at $z_0$. This leaves:

$$\partial_t N(t,TB) \sim -\,c(t)\,\rho(z_0,t,TB). \tag{59}$$

From Eqns. (44) and (45), we can readily derive an approximate expression for this leakage using the quasi-steady solutions for $\rho(z_0,t,TB)$. Using the fact that $s(z_0,t) = 0$ by definition, and plugging in Eqn. (44) gives:

$$\partial_t N(t,TB) \sim -\,\epsilon \frac{c(t)^2}{\langle \chi(TB) - \mu(TB) | z_0, t\rangle} P(TB|z_0,t)\,N(t). \tag{60}$$

With the arguments in the previous section, the presence of low-performing phenotypes in the migrating group dramatically reduces the density of high performers at the back, thus dramatically reducing their leakage flux.

Finally, the total leakage flux is:

$$\partial_t N(t) = \int \partial_t N(t,TB)\, dTB = -c(t)\, \rho(z_0,t) \sim -\, \epsilon \frac{c(t)^2}{\langle \chi(TB) - \mu(TB)|z_0,t\rangle}\, N(t). \tag{61}$$

### Growth model

When a cell with phenotype $TB$ divides, what are the phenotypes of its daughter cells? We model the probability of a cell dividing in time $dt$ during exponential growth as a Poisson process. One could then construct a master equation with microscopic transitions that are proportional to the rates at which mother cells divide and the relative probabilities of producing two daughter cells with some phenotypes. Ultimately, however, only the average net growth matrix appears in the PDE for cell density through an additional term on the right-hand side:

$$\int R(TB,TB',z)\, \rho(z,t,TB')\, dTB'. \tag{62}$$

For a discrete set of phenotypes, element $R_{ij}(z) = R\big(TB_i,TB_j,z\big)$ is the average net rate at which cells of phenotype $TB_j$ produce cells of phenotype $TB_i$. For example, dropping the $z$-dependence for now, on the diagonal, $R_{ii} = r_i\,\big(2\, p_{ii|i} + \sum_j p_{ij|i} + \sum_j p_{ji|i} - 1\big)$ has four contributions. Here, $r_i$ is the rate at which phenotype $TB_i$ undergoes divisions. The first term is the probability that both daughter cells have the same phenotype as the mother, with the factor of 2 accounting for two cells being produced. The second and third terms are the probability that only one daughter cell has the same phenotype as the mother. The last term accounts for removal of the mother cell upon division. This formulation allows for cases where the two daughter cells are distinguishable, such as the “old pole” cell and the “new pole”, and could have different, possibly correlated, statistics. The other entries of the matrix $R$ have terms that include the possibility that division of a cell with phenotype $TB_j$ produces two cells with phenotype $TB_i$. The sum of elements in each column of $R$ is $r_j$, the division rate of phenotype $TB_j$.

The structure of $R$ that arises from the microscopic transitions described above places constraints on its elements. The diagonal elements can be less than zero, but the remaining elements must be non-negative because division of one phenotype cannot cause removal of another phenotype. Additionally, the sum of all elements in a column, excluding the diagonal element, must be less than or equal to $2\, r_j$, with equality occurring only when phenotype $j$ never produces offspring with the same phenotype as itself.

In general, the growth matrix $R$ depends on position $z$ through its dependence on the local concentration of nutrients. This can be simplified by assuming that the growth nutrients are abundant and distinct from the attractant, making the growth matrix independent of position: $R = R(TB,TB')$. Furthermore, we assume that all phenotypes grow exponentially with the same maximal rate, $r_j = r$. Additionally, the dynamics of cell density do not distinguish which daughter cell is which, and therefore the joint distribution of daughter phenotypes is symmetric. In this case, even if the daughter cell phenotypes are correlated, $R(TB,TB')$ only depends on the marginal distributions of each daughter cell’s phenotype. To see this, we write

$$\begin{aligned} R_{ij} = r\, \Big(&2\, P\big(TB_1 = TB_i, TB_2 = TB_i \big| TB_j\big) + P\big(TB_1 = TB_i, TB_2 \neq TB_i \big| TB_j\big) \\ &+ P\big(TB_1 \neq TB_i, TB_2 = TB_i \big| TB_j\big) - \delta\big(TB_i, TB_j\big)\Big) \end{aligned} \tag{63}$$

with $\delta\big(TB_i, TB_j\big) = 1$ if $TB_i = TB_j$ and $\delta\big(TB_i, TB_j\big) = 0$ otherwise. If the distribution of the two daughter phenotypes is symmetric, then $P\big(TB_1 = TB_i, TB_2 \neq TB_i \big| TB_j\big) = P\big(TB_1 \neq TB_i, TB_2 = TB_i \big| TB_j\big)$. Then $R_{ij}$ simplifies to:

$$R_{ij} = r\,\Big(2\,\Big(P\big(TB_1 = TB_i, TB_2 = TB_i \big| TB_j\big) + P\big(TB_1 = TB_i, TB_2 \neq TB_i \big| TB_j\big)\Big) - \delta\big(TB_i, TB_j\big)\Big). \tag{64}$$

But $P\big(TB_1 = TB_i, TB_2 = TB_i \big| TB_j\big) + P\big(TB_1 = TB_i, TB_2 \neq TB_i \big| TB_j\big)$ is the marginalization of $P\big(TB_1, TB_2 \big| TB_j\big)$ over $TB_2$. Due to the symmetry between $TB_1$ and $TB_2$, this leaves:

$$R_{ij} = r\,\Big(2\,P\big(TB_i \big| TB_j\big) - \delta\big(TB_i, TB_j\big)\Big). \tag{65}$$

Symmetry of the joint distribution of the two daughter cell phenotypes makes their marginal distributions the same. As a result, even if the phenotypes of daughter cells are correlated, only their marginal distributions appear in $R(TB, TB')$.

The PDE for cell density in the moving reference frame (Eqn. (10)) becomes:

$$\begin{aligned}\partial_t \rho(z,t,TB) &- c(t)\,\partial_z \rho(z,t,TB) \\ &= -\partial_z\left(\chi(TB)\,\partial_z \log\left(\epsilon + \frac{s(z,t)}{s_\infty}\right)\rho(z,t,TB) - \mu(TB)\partial_z\rho(z,t,TB)\right) \\ &+ r\int \big(2\,P(TB|TB') - \delta(TB - TB')\big)\,\rho(z,t,TB')\,dTB'.\end{aligned} \tag{66}$$

Next, we need a model for $P(TB|TB')$. We expect that daughter cells should have phenotypes similar to their mothers, with some variation. Possibly the simplest model of growth with local inheritance of phenotype is an autoregressive (AR) process, where the phenotypes of daughter cells are Gaussian-distributed with means that revert back to the batch culture mean phenotype. This model is more appropriate for phenotypes defined over $(-\infty, \infty)$, rather than the finite region $[0,1]$ on which $TB$ is defined. Therefore, we consider an AR-type growth-inheritance model for $F$ (7), defined by:

$$TB = \frac{1}{1 + \exp(F)}, \qquad F = \log\left(\frac{1 - TB}{TB}\right). \tag{67}$$

$F$ can be thought of as an effective free energy difference between the run and tumble states. As $F \to \infty$, $TB \to 0$; as $F \to -\infty$, $TB \to 1$. Since the mapping from $F$ to $TB$ is sigmoidal, cells with $TB$ approaching zero or one are increasingly difficult to generate.

The AR model for the distribution of offspring phenotypes $F$ given mother's phenotype $F'$ has 3 parameters: the mean $\langle F\rangle$ and variance $\sigma^2$ of $F$ in the batch culture distribution, and $\phi$ the correlation between mother and daughter phenotypes. In this model, the offspring distribution is:

$$P(F|F') = \mathcal{N}\big((1-\phi)\,\langle F\rangle + \phi\, F', \sigma^2(1-\phi^2)\big) \tag{68}$$

where $\mathcal{N}(\mu, \sigma^2)$ is a Gaussian distribution with mean $\mu$ and variance $\sigma^2$. Again, since we assume the two daughter cell phenotype statistics are symmetric, we only need to specify their marginal distribution $P(F|F')$. As $\phi \to 1$, daughter cells become identical to their mothers, and the population never reaches a steady state composition in batch culture. When $\phi = 0$, daughter cells are sampled directly from the batch culture distribution. The relaxation time of the population composition in batch culture is found from the first nonzero eigenvalue of $P(F|F') - \delta(F - F')$, which in this model is $\lambda_1 = -(1-\phi)$, with corresponding eigenfunction $\partial_F P(F|F')$. The relaxation time of the population composition in batch culture is then $\tau_r = -\frac{1}{2\,r\,\lambda_1} = \frac{1}{2\,r\,(1-\phi)}$. Thus, the correlation between mother and daughter phenotypes controls the time scale on which the population's phenotype composition relaxes to steady state in batch culture.

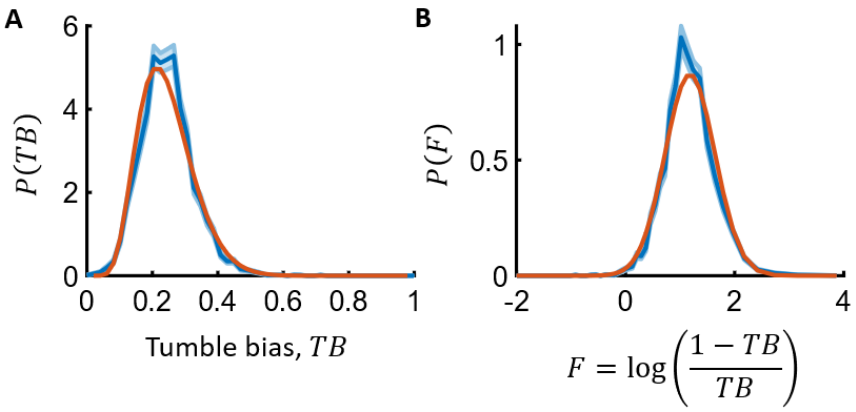

**Figure S3: Wild type distribution of phenotypes $TB$ in batch culture.** The tumble bias distribution $P(TB)$ of wild-type *E. coli* cells is well-described by a Gaussian distribution of the transformed phenotype $F = \log\left(\frac{1-TB}{TB}\right)$. Tumble bias distributions were measured by tracking RP437 cells (blue, left; shading throughout is standard error found by bootstrapping). Cells picked from a -80°C freezer stock were grown in 1X M9 salts + 0.1% w/v casamino acids + 0.4% v/v glycerol + 1 mM MgSO4 + 0.05% w/v PVP-40 (1) at 30°C overnight, and then diluted 100X in the same media the following morning, and harvested at OD600 = 0.25. The cells were then washed three times in chemotaxis buffer (1X M9 salts + 10 µM methionine + 100 µM EDTA + 0.05% w/v PVP-40) (1), injected into a 100 um-deep microfluidic chamber, and tracked with a 4X objective by phase contrast imaging at 20 frames per second. Three biological replicates were performed, and only trajectories longer than 10 seconds were kept, resulting in 5,825 trajectories with mean duration of 20 seconds. Tumbles were detected using angular speed and swimming speed variations, as described in refs (17,30). From $P(TB)$, the distribution $P(F)$ was computed using a change of variables formula (blue, right). $P(F)$ was approximated by a Gaussian distribution with the same mean and variance (orange, right). This distribution was then transformed to a distribution of tumble bias by inverting the change of variable formula (left, orange). The mean of $P(F)$ was 1.2 and the standard deviation was 0.46. Simulations throughout used a different mean phenotype (explained in Fig. S5 caption), but the same amount of variation.

### Steady state migration speed

Steady state migration is reached when the number of cells of each phenotype produced per unit time is equal to the number lost per unit time. The production of cells in the group is integral of the growth term over the width of the wave, i.e. from $z = z_0$ to $z \to \infty$. Since we assumed the growth matrix is independent of position, this reduces to

$$[\partial_t N(t,TB)]_{growth} = \int_{z_0}^{\infty} \int R(TB,TB')\, \rho(z,t,TB')\, dTB' \ dz \tag{69}$$

$$= \int R(TB,TB')\, N(t,TB')\, dTB'. \tag{70}$$

Then, the time evolution of $N(t,TB)$ in the wave during travel has contributions from growth and leakage:

$$\frac{dN(t,TB)}{dt} = [\partial_t N(t,TB)]_{growth} + [\partial_t N(t,TB)]_{leakage}. \tag{71}$$

Since the time scale of growth, $1/r$, is typically slow compared to the time scale of wave migration, $\langle \chi(TB) - \mu(TB) \rangle / c(t)^2$ (21), the quasi-stationary expression for $\rho(z,t,TB)$ and the leakage expression derived above are expected to be a good approximation in the presence of growth. Using these, we get:

$$\frac{dN(t,TB)}{dt} = \int R(TB,TB')\, N(t,TB')\, dTB' - \epsilon \frac{c(t)^2}{\langle \chi(TB) - \mu(TB) | z_0, t \rangle} \frac{P(TB|z_0,t)}{P(TB,t)}\, N(t,TB) \tag{72}$$

$$= \int r \left( 2\, P(TB|TB') - \delta(TB - TB') \right) N(t,TB')\, dTB' - \epsilon \frac{c(t)^2}{\langle \chi(TB) - \mu(TB) | z_0, t \rangle} \frac{P(TB|z_0,t)}{P(TB,t)}\, N(t,TB) \tag{73}$$

The total number of cells is the integral of the above expression over phenotypes $TB$:

$$\frac{dN(t)}{dt} = \int 2\, r \int P(TB|TB')\, N(t,TB')\, dTB' \ dTB - r\, N(t) - \epsilon \frac{c(t)^2}{\langle \chi(TB) - \mu(TB) | z_0, t \rangle}\, N(t). \tag{74}$$

Integrating the first term with respect to $TB$ first results in the following simple expressions:

$$\frac{dN(t)}{dt} = 2\, r \int N(t,TB')\, dTB' - r\, N(t) - \epsilon \frac{c(t)^2}{\langle \chi(TB) - \mu(TB) | z_0, t \rangle}\, N(t) \tag{75}$$

$$= r\, N(t) - \epsilon \frac{c(t)^2}{\langle \chi(TB) - \mu(TB) | z_0, t \rangle}\, N(t). \tag{76}$$

At steady state, growth and leakage balance

$$\frac{dN(t)}{dt} = 0 \to r\, N_{ss} = \epsilon \frac{c_{ss}^2}{\langle \chi(TB) - \mu(TB) | z_0, ss \rangle}\, N_{ss}, \tag{77}$$

and solving for $c_{ss}$:

$$c_{ss} = \left(\frac{r}{\epsilon}\langle \chi(TB) - \mu(TB) | z_0, ss\rangle\right)^{1/2}. \tag{78}$$

We find that the steady state traveling speed is set by the average performance of phenotypes at the back of the wave at steady state, $\langle \chi(TB) - \mu(TB) | z_0, ss\rangle$. Although this quantity is difficult to predict, it is clear that if the composition of the migrating group shifts such that minimum performance in higher, then the wave speed at steady state will be higher.

This theoretical expression is consistent with recent measurements (9) for the dependence of wave speed on growth rate $r$ and its lack of dependence on the consumption rate of attractant $k$ (assuming that the wave composition was similar across those experiments). Recent theory that treated all cells as identical and considered attractant diffusion found the same dependence on $r$, $\epsilon$, and $k$ (21). We extend the expression in Eqn. (78) to account for attractant diffusion in the following section, **Steady state migration speed with attractant diffusion**.

Finally, as recently found for migrating populations consisting of a single phenotype, the density of cells left behind diverse migrating waves at steady state also has a simple form. At steady state, the flux of cells off the back is balanced by the total flux of cells produced by growth:

$$c_{ss}\, \rho_{ss}(z_0, TB) = r\, N_{ss} \rightarrow \rho_{ss}(z_0, TB) = r\frac{N_{ss}}{c_{ss}} = r\frac{s_\infty}{k}, \tag{79}$$

where we used $c_{ss} = k\, N_{ss} \,/\, s_\infty$. Surprisingly, the total density of cells left behind is independent of the steady state phenotype composition of cells traveling when growth nutrients are in excess within the group, and therefore is also independent of phenotypic inheritance and independent of which physical environment the cells are migrating through.

### Steady state migration speed with attractant diffusion

As noted recently (21), attractant diffusion can have significant effects on the migrating wave when the attractant diffusivity $D_s$ approaches or exceeds $\chi(TB) - \mu(TB)$ among the cells traveling. This can occur in porous environments, where all phenotypes' chemotactic coefficients and diffusivities are lower than in liquid, but the attractant diffusivity is essentially unchanged. The steady state ODEs we will consider are slightly different from those in earlier sections. In addition to keeping attractant diffusion, the form of the consumption term will be that used in (21):

$$-c_{ss}\, \partial_z \left(\frac{s}{s_\infty}\right) = D_s\, \partial_{zz}\left(\frac{s}{s_\infty}\right) - \frac{k}{s_\infty}\left(\frac{\frac{s}{s_\infty}}{\frac{s}{s_\infty} + \epsilon}\right)\rho. \tag{80}$$

Throughout, the $z$-dependence of $\rho$ and $s$ are understood, but not shown, $\rho$ is total cell density, $\rho(TB)$ is the density of cells with phenotype $TB$, and $c_{ss}$ is the steady state migration speed. Importantly, this consumption term has a Monod-like form with a half-maximum consumption constant $K_m$ that we set equal to $K_i$, the chemotactic sensitivity for the attractant. Thus, $K_m/s_\infty = K_i/s_\infty = \epsilon$.

Our approach in this section will closely follow that of (21), but with non-genetic diversity of swimming phenotypes. We focus on the back region of the migrating group, where chemotaxis dominates over cell diffusion, and make the ansatz that the attractant has the following form in the chemotaxis region:

$$\frac{s}{s_\infty} = A \exp\left(\int \lambda(z)\, dz\right). \tag{81}$$

Plugging the attractant ansatz into the attractant ODE, Eqn. (79) gives:

$$c_{ss}\, \lambda(z) \frac{s}{s_\infty} + D_s\, (\partial_z \lambda(z) + \lambda(z)^2) \frac{s}{s_\infty} = \frac{k}{s_\infty} \frac{\rho}{\left(\epsilon + \frac{s}{s_\infty}\right)} \frac{s}{s_\infty} \tag{82}$$

$$c_{ss}\, \lambda(z) \frac{s}{s_\infty} + D_s\, \lambda(z)\ \left(\partial_z \log\big(\lambda(z)\big) + \lambda(z)\right) \frac{s}{s_\infty} = \frac{k}{s_\infty} \frac{\rho}{\left(\epsilon + \frac{s}{s_\infty}\right)} \frac{s}{s_\infty}. \tag{83}$$

Next, we argue that $\lambda(z) \gg \partial_z \log\big(\lambda(z)\big)$. Later, we'll see that $\lambda(z) = \frac{c_{ss}}{\langle \chi - \mu | z \rangle}$, making $\partial_z \log\big(\lambda(z)\big) = -\partial_z \log(\langle \chi | z \rangle - \langle \mu | z \rangle)$. With this, the assumption that $\lambda(z) \gg \partial_z \log\big(\lambda(z)\big)$ means that although chemotaxis abilities in the group vary with position, but they vary on a length scale that is longer than the width of group itself, $\sim 1/\lambda(z)$. If the opposite were true, i.e. $\lambda(z) \ll \partial_z \log\big(\lambda(z)\big)$, then cells throughout the group would experience almost the same gradient. In this case, there would be no compensatory mechanism keeping them traveling together, so a stable traveling wave cannot form in this regime. We also argue that it is not the case $\lambda(z) \sim \partial_z \log\big(\lambda(z)\big)$, because empirically in simulations, chemotaxis abilities in the group do not change by orders of magnitude in the region where chemotaxis dominates. Thus, we take $\lambda(z) \gg \partial_z \log\big(\lambda(z)\big)$, and we simplify the ODE above to:

$$c_{ss}\, \lambda(z) + D_s\, \lambda(z)^2 = \frac{k}{s_\infty} \frac{\rho}{\left(\epsilon + \frac{s}{s_\infty}\right)}. \tag{84}$$

We'll return to this equation later.

Next, we consider the ODE for $\rho(TB)$, reproduced here:

$$-c_{ss}\, \partial_z \rho(TB) = -\partial_z \left( \chi(TB) \frac{\rho(TB)}{\left(\epsilon + \frac{s}{s_\infty}\right)}\, \partial_z \left(\frac{s}{s_\infty}\right) - \mu(TB)\, \partial_z \rho(TB) \right) + r \int R(TB|TB')\, \rho(TB')\, dTB'. \tag{85}$$

We will immediately integrate this equation over phenotypes $TB$.

$$-c_{ss}\,\partial_z\rho = -\partial_z\left(\int \chi(TB)\,\rho(TB)\,dTB\,\frac{\partial_z\left(\frac{s}{s_\infty}\right)}{\left(\epsilon+\frac{s}{s_\infty}\right)} - \int \mu(TB)\,\partial_z\rho(TB)\,dTB\right) + r\int\int R(TB|TB')\,\rho(TB')\,dTB'\,dTB. \tag{86}$$

To simplify this equation, we first note that $P(TB|z) = \rho(TB)/\rho$. With this,

$$\int \chi(TB)\,\rho(TB)\,dTB = \rho\int \chi(TB)\frac{\rho(TB)}{\rho}\,dTB = \rho\,\langle\chi|z\rangle. \tag{87}$$

Furthermore, we can simplify the last term by changing the order of integration with respect to $TB$ and $TB'$, and using the fact that $\int R(TB|TB')\,dTB = 1$ for all $TB'$:

$$\int\left(\int R(TB|TB')\,\rho(TB')\,dTB'\right)dTB \tag{88}$$

$$= \int\left(\int R(TB|TB')\,dTB\right)\rho(TB')\,dTB' = \int \rho(TB')\,dTB' = \rho. \tag{89}$$

Using these simplifications, Eqn. (86) becomes:

$$0 = -\partial_z\left(\langle\chi|z\rangle\frac{\rho}{\epsilon+\frac{s}{s_\infty}}\,\partial_z\left(\frac{s}{s_\infty}\right) - c_{ss}\,\rho - \int \mu(TB)\,\partial_z\rho(TB)\,dTB\right) + r\,\rho. \tag{90}$$

To simplify the term $\int \mu(TB)\,\partial_z\rho(TB)\,dTB$, note that:

$$\partial_z P(TB|z) = \partial_z\frac{\rho(TB)}{\rho} = \frac{1}{\rho}\partial_z\rho(TB) - \frac{\rho(TB)}{\rho^2}\partial_z\rho = \frac{1}{\rho}\partial_z\rho(TB) - \frac{1}{\rho}P(TB|z)\,\partial_z\rho \tag{91}$$

And therefore

$$\partial_z\rho(TB) = \rho\,\partial_z P(TB|z) + P(TB|z)\,\partial_z\rho. \tag{92}$$

With this, we get

$$\int \mu(TB)\,\partial_z\rho(TB)\,dTB = \int \mu(TB)\,(\rho\,\partial_z P(TB|z) + P(TB|z)\,\partial_z\rho)\,dTB \tag{93}$$

$$= \rho\int \mu(TB)\,\partial_z P(TB|z)\,dTB + \langle\mu(TB)|z\rangle\,\partial_z\rho \tag{94}$$

$$= \rho\,\partial_z\langle\mu|z\rangle + \langle\mu|z\rangle\,\partial_z\rho \tag{95}$$

Plugging this into Eqn. (90):

$$0 = -\partial_z \left( \langle \chi | z \rangle \frac{\rho}{\epsilon + \frac{s}{s_\infty}} \, \partial_z \left( \frac{s}{s_\infty} \right) - c_{ss} \, \rho - (\rho \, \partial_z \langle \mu | z \rangle + \langle \mu | z \rangle \, \partial_z \rho) \right) + r \, \rho \tag{96}$$

$$0 = -\partial_z \left( \langle \chi | z \rangle \frac{\rho}{\epsilon + \frac{s}{s_\infty}} \, \partial_z \left( \frac{s}{s_\infty} \right) - c_{ss} \, \rho - \langle \mu | z \rangle \, \rho \, (\partial_z \log(\langle \mu | z \rangle) + \partial_z \log(\rho)) \right) + r \, \rho \tag{97}$$

Plugging in the attractant ansatz:

$$0 = -\partial_z \left( \langle \chi | z \rangle \frac{\rho}{\epsilon + \frac{s}{s_\infty}} \, \lambda(z) \, \frac{s}{s_\infty} - c_{ss} \, \rho - \langle \mu | z \rangle \, \rho \, (\partial_z \log(\langle \mu | z \rangle) + \partial_z \log(\rho)) \right) + r \, \rho. \tag{98}$$

Next, we make the following ansatz for total cell density: $\rho = \beta(z) \left( \epsilon + \frac{s}{s_\infty} \right)$. With this ansatz, $\partial_z \log(\rho) = \partial_z \log\big(\beta(z)\big) + \frac{1}{\left( \epsilon + \frac{s}{s_\infty} \right)} \lambda(z) \frac{s}{s_\infty}$. Later, we will see that $\partial_z \log\big(\beta(z)\big) \ll \lambda(z)$, and therefore:

$$\partial_z \log(\rho) \sim \frac{1}{\left( \epsilon + \frac{s}{s_\infty} \right)} \lambda(z) \frac{s}{s_\infty} = O\big(\lambda(z)\big). \tag{99}$$

We anticipate, from our derivations in earlier sections, that the gradient steepness $\lambda(z)$ will be:

$$\lambda(z) = \frac{c_{ss}}{\langle \chi - \mu | z \rangle}. \tag{100}$$

Furthermore, if $\chi/\mu$ is constant among all phenotypes, which we assumed earlier, we have:

$$\partial_z \log(\langle \mu | z \rangle) = \partial_z \log \left( \frac{(\alpha - 1)}{c_{ss}} \langle \mu | z \rangle \right) \tag{101}$$

$$= \partial_z \log \left( \frac{\langle \chi - \mu | z \rangle}{c_{ss}} \right) \tag{102}$$

$$= -\partial_z \log\big(\lambda(z)\big). \tag{103}$$

With our claim that $\partial_z \log\big(\lambda(z)\big) \ll \lambda(z)$, we then have $\partial_z \log(\langle \mu | z \rangle) \ll \lambda(z)$, and therefore $\partial_z \log(\langle \mu | z \rangle) \ll \partial_z \log(\rho)$. As a result, we drop $\partial_z \log(\langle \mu | z \rangle)$ from the ODE for $\rho$:

$$0 = -\partial_z \left( \langle \chi | z \rangle \frac{\rho}{\epsilon + \frac{s}{s_\infty}} \, \lambda(z) \frac{s}{s_\infty} - c_{ss} \, \rho - \langle \mu | z \rangle \, \rho \, \partial_z \log(\rho) \right) + r \, \rho. \tag{104}$$

Plugging in the ansatz $\rho = \beta(z) \left( \epsilon + \frac{s}{s_\infty} \right)$ and $\partial_z \log(\rho) \sim \frac{1}{\left( \epsilon + \frac{s}{s_\infty} \right)} \lambda(z) \frac{s}{s_\infty}$:

$$0 = -\partial_z \left( \langle \chi|z \rangle \, \beta(z) \, \lambda(z) \frac{s}{s_\infty} - c_{ss} \, \beta(z) \left( \epsilon + \frac{s}{s_\infty} \right) - \langle \mu|z \rangle \, \beta(z) \, \lambda(z) \frac{s}{s_\infty} \right) + r \, \beta(z) \left( \epsilon + \frac{s}{s_\infty} \right) \quad (105)$$

Rearranging terms:

$$0 = -\partial_z \left( \beta(z) \left( (\langle \chi|z \rangle - \langle \mu|z \rangle) \, \lambda(z) \frac{s}{s_\infty} - c_{ss} \, \left( \epsilon + \frac{s}{s_\infty} \right) \right) \right) + r \, \beta(z) \left( \epsilon + \frac{s}{s_\infty} \right) \quad (106)$$

Next, we plug in the expected form of the gradient steepness, $\lambda(z) = \frac{c}{\langle \chi|z \rangle - \langle \mu|z \rangle}$:

$$0 = -\partial_z \left( \beta(z) \left( c_{ss} \frac{s}{s_\infty} - c_{ss} \, \left( \epsilon + \frac{s}{s_\infty} \right) \right) \right) + r \, \beta(z) \left( \epsilon + \frac{s}{s_\infty} \right) \quad (107)$$

$$\partial_z \beta(z) = -\frac{r}{c_{ss} \, \epsilon} \, \beta(z) \left( \epsilon + \frac{s}{s_\infty} \right) \quad (108)$$

$$\partial_z \log(\beta(z)) = -\frac{r}{c_{ss} \, \epsilon} \left( \epsilon + \frac{s}{s_\infty} \right) \quad (109)$$

Eqn. (109) above indicates that $\partial_z \log(\beta(z)) \ll \lambda(z)$ specifically in the region where $\frac{s}{s_\infty} \sim \epsilon$, because in that region $\partial_z \log(\beta(z)) \sim \frac{r}{c_{ss}} \ll \lambda(z)$ (as in (21)). This is enough for our purposes, because this is the region of interest for leakage. This also means that our guess $\lambda(z) = \frac{c_{ss}}{\langle \chi|z \rangle - \langle \mu|z \rangle}$ is valid in that region – there are no contradictions.

Next, we integrate the ODE for total cell density $\rho$ Eqn. (96) over space from $z$ to $z \to \infty$. Although the ansatz is no longer valid as $z \to \infty$, it is only a local approximation of the actual function $\rho$. Therefore, we apply the boundary conditions in Eqn. (12) to get:

$$c_{ss} \, \rho = \langle \chi|z \rangle \frac{\rho}{\epsilon + \frac{s}{s_\infty}} \, \partial_z \frac{s}{s_\infty} - (\rho \, \partial_z \langle \mu|z \rangle + \langle \mu|z \rangle \, \partial_z \rho) + r \, N(z). \quad (110)$$

All of the calculations that we made inside the $z$-derivative in the section above still apply. With them, we get:

$$0 = \beta(z) \left( (\langle \chi|z \rangle - \langle \mu|z \rangle) \, \lambda(z) \frac{s}{s_\infty} - c_{ss} \, \left( \epsilon + \frac{s}{s_\infty} \right) \right) + r \, N(z), \quad (111)$$

where $N(z)$ is the cumulative number of cells in the traveling group ahead of position $z$.

Substituting for $\lambda(z)$:

$$0 = \beta(z) \left( c_{ss} \frac{s}{s_\infty} - c_{ss} \, \left( \epsilon + \frac{s}{s_\infty} \right) \right) + r \, N(z) \quad (112)$$

$$\beta(z) = \frac{r}{c_{ss} \, \epsilon} \, N(z) \quad (113)$$

Again, this is valid at the back, where $\frac{s}{s_\infty} \sim \epsilon$. Evaluating both sides at the back of the wave $z = z_0$ and use $N(z_0) = N$, the total number of cells traveling:

$$\beta(z_0) \sim \frac{r}{c_{ss}\,\epsilon} N. \tag{114}$$

Here, we define $z_0$ as the location where $\frac{s(z_0)}{s_\infty} = \epsilon$. We will see below that this captures the total number of cells to leading order.

Next, we integrate the attractant ODE from $z = z_0$ to $z \to \infty$ and apply the boundary conditions in Eqn. (12) again:

$$-c_{ss}\,(1-\epsilon) = D_s \int_{z_0}^{\infty} \left( \partial_{zz}\left(\frac{s}{s_\infty}\right) - \frac{k}{s_\infty} \left( \frac{\frac{s}{s_\infty}}{\frac{s}{s_\infty} + \epsilon} \right) \rho \right) dz. \tag{115}$$

Using the boundary conditions, the ansatz for the attractant, and the definition $\frac{s(z_0)}{s_\infty} = \epsilon$, we can simplify the first term on the right-hand side:

$$-c_{ss}\,(1-\epsilon) = -D_s\,\lambda(z_0)\,\epsilon - \int_{z_0}^{\infty} \frac{k}{s_\infty} \left( \frac{\frac{s}{s_\infty}}{\frac{s}{s_\infty} + \epsilon} \right) \rho\, dz. \tag{116}$$

The last term can be split into two:

$$-c_{ss}\,(1-\epsilon) = -D_s\,\lambda(z_0)\,\epsilon - \left( \int_{z_0}^{\infty} \frac{k}{s_\infty}\, \rho\, dz - \int_{z_0}^{\infty} \frac{k}{s_\infty} \left( \frac{\epsilon}{\frac{s}{s_\infty} + \epsilon} \right) \rho\, dz \right) \tag{117}$$

$$-c_{ss}\,(1-\epsilon) = -D_s\,\lambda(z_0)\,\epsilon - \left( \frac{k}{s_\infty} N - \int_{z_0}^{\infty} \frac{k}{s_\infty} \left( \frac{\epsilon}{\frac{s}{s_\infty} + \epsilon} \right) \rho\, dz \right) \tag{118}$$

The last term here is small (i.e. $O(\epsilon)$) because in regions where $\frac{s}{s_\infty} \sim O(\epsilon)$, we also have $\rho \sim O(\epsilon)$, and in regions where $\frac{s}{s_\infty} \gg \epsilon$, the term in the parenthesis is $O(\epsilon)$. Keeping only leading-order terms, we get:

$$c_{ss} = \frac{k}{s_\infty} N, \tag{119}$$

as in Keller and Segel's analysis.

Plugging this into Eqn. (114), we eliminate $N$ and solve for $\beta(z_0)$:

$$\beta(z_0) = \frac{r}{\epsilon} \frac{s_\infty}{k}. \tag{120}$$

Finally, we return to the ODE for the attractant $s$, Eqn. (84). Plugging in the ansatz for $\rho$:

$$c_{ss}\, \lambda(z) + D_s\, \lambda(z)^2 = \frac{k}{s_\infty} \beta(z) \tag{121}$$

Evaluating both sides at $z = z_0$ and plugging in expression for $\lambda(z_0)$:

$$\frac{c_{ss}{}^2}{\langle \chi - \mu | z_0 \rangle} + D_s \left( \frac{c_{ss}}{\langle \chi - \mu | z_0 \rangle} \right)^2 = \frac{r}{\epsilon}. \tag{122}$$

Solving for the steady state migration speed $c_{ss}$, we find that

$$c_{ss} = \left( \frac{r}{\epsilon} \frac{\langle \chi - \mu | z_0 \rangle^2}{\langle \chi - \mu | z_0 \rangle + D_s} \right)^{1/2}. \tag{123}$$

Note that both Eqn. (31) and Eqn. (119) show that the migration speed $c$ at any given time is proportional to the number of cells traveling $N$, but they differ in how the back of the group $z_0$ is defined, and therefore differ in terms of which cells count towards $N$ because $N \equiv \int_{z_0}^{\infty} \rho\, dz$. When $K_m \to 0$, the back of the group $z_0$ is defined by $s(z_0) = 0$, and in simulations this definition of the back does indeed make $N = c\, s_\infty / k$. When $K_m = K_i$, instead defining $z_0$ such that $s(z_0) = K_m = K_i$ makes $N = c\, s_\infty / k$. For more general $K_m$ and $K_i$, we find empirically in simulations that $N = c\, s_\infty / k$ results when the back of the group $z_0$ is defined by $s(z_0) = \sqrt{K_m\, K_i}$. This captures both of the case we considered analytically here, as well as the Keller-Segel case of $K_i \to 0$ (and $z_0 \to -\infty$).

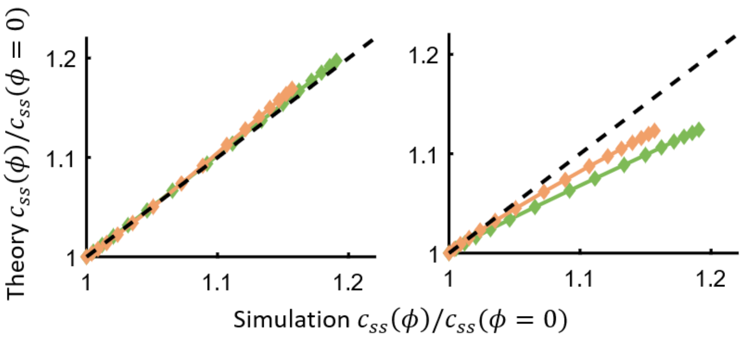

**Figure S4: Steady state speed predictions.** Top: Predicted steady state migration speed versus that in simulations, for varying value of mother-daughter inheritance $\phi$ and normalized by the value at $\phi = 0$ (predictions made by taking $\langle \chi - \mu | z_0 \rangle$ from simulations and using Eqn. 11 of the main text, or Eqn. (123) of the SI; reproduced from Fig. 3D). Bottom: the same as top, but where the predictions use the mean

$\chi - \mu$ over all cells traveling Eqn. (123), rather than $\langle \chi - \mu | z_0 \rangle$, indicating that the former cannot accurately describe steady state migration speed.

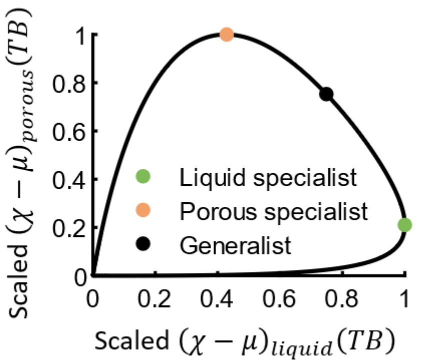

**Figure S5: Trade-offs in chemotaxis performance between liquid and porous environments.** Plotted is $\chi(TB) - \mu(TB)$ in porous media versus that in liquid, where each is normalized by its maximum value across phenotypes $TB$. The green dot indicates the liquid specialist phenotype. Among populations containing just one phenotype, it would travel fastest at steady state in liquid if it had this phenotype ($TB \sim 0.02$ in our model). Likewise, the orange dot indicates the porous specialist ($TB \sim 0.31$ in our model). The black dot indicates the generalist phenotype, which has the highest minimum value of normalized $\chi(TB) - \mu(TB)$ across the two environments ($TB \sim 0.12$ in our model). A population consisting only of this phenotype would likewise have the highest minimum steady state migration speed across environments, among all phenotypes. Throughout, in simulations with diversity, the most abundant phenotype was chosen to be the generalist, and the amount of variation was chosen to be the same as the wild type population in Fig. S3.

### Total population growth

As shown in experiments, total population growth, including cells behind the wave, increases as the migration speed increases (9). This is true for a variety of situations. First we consider the case that both cells in the wave and behind it grow with rate $r$. Since the density of cells left behind the wave is always constant at steady state, Eqn. (79), we can write the density of cells behind the wave as

$$\rho(x,t) = \rho_{leak} \exp(r(t - t_{leak}(x)) = r\frac{S_\infty}{k} \exp\big(r(t - t_{leak}(x)\big), \text{for } t - t_{leak}(x) > 0. \quad (124)$$

where $t_{leak}(x)$ is the time at which cells were left behind the wave at position $x$. The current location of the back of the group is given by $x = c_{ss}\, t$ (up to a constant shift), and therefore $t_{leak}(x) = x/c_{ss}$. Then, the total population size $N_{tot}(t)$ after traveling for time $t$ at steady state is the sum over cells behind the group plus those in the group, $N_{ss} = c_{ss}\frac{S_\infty}{k}$:

$$N_{tot}(t) = \int_0^x \rho(x,t)\,dx + N_{ss} \tag{125}$$

$$= \int_0^{c_{ss}\,t} r\frac{s_\infty}{k}\exp\left(r\left(t-\frac{x}{c_{ss}}\right)\right)dx + c_{ss}\frac{s_\infty}{k} \tag{126}$$

$$= r\frac{s_\infty}{k}e^{r\,t}\left(-\frac{c_{ss}}{r}(e^{-rt}-1)\right) + c_{ss}\frac{s_\infty}{k} \tag{127}$$

$$= c_{ss}\frac{s_\infty}{k}\,e^{r\,t}. \tag{128}$$

Importantly, directed range expansion in the presence of abundant nutrients enables the population to maintain exponential growth.

Even if just the cells in the migrating group grow with rate $r$, cells are produced at the same rate as they are leaked behind, so the population size grows as

$$N_{tot}(t) = c_{ss}\,\rho_{leak}\,t = c_{ss}\,r\,\frac{s_\infty}{k}\,t. \tag{129}$$

In both cases, faster migration leads to faster total population growth.

Population expansion driven by growth and cell diffusion, as in Fisher waves, grows the population size like:

$$N_{tot}(t) = c_{ss}\,\rho_{max}\,t = \sqrt{r\,\mu}\,\rho_{max}\,t, \tag{130}$$

where $\rho_{max}$ is the carrying capacity, $c_{ss} = \sqrt{r\,\mu}$ is the Fisher wave speed, $r$ is again the growth rate, and we've taken the diffusivity of all cells to be $\mu$.
